# Cardiolipin coordinates inflammatory metabolic reprogramming through regulation of Complex II assembly and stability

**DOI:** 10.1101/2022.07.27.501734

**Authors:** Mack B. Reynolds, Hanna S. Hong, Britton C Michmerhuizen, Anna-Lisa E. Lawrence, Li Zhang, Jason S. Knight, Costas A. Lyssiotis, Basel H. Abuaita, Mary X. O’Riordan

## Abstract

Macrophage metabolic plasticity enables repurposing of electron transport from energy generation to inflammation and host defense. Altered Respiratory Complex II function has been implicated in cancer, diabetes and inflammation but regulatory mechanisms are incompletely understood. Here we show that macrophage inflammatory activation triggers Complex II disassembly and succinate dehydrogenase-B subunit loss through sequestration and mitophagy. Mitochondrial fission was required for lipopolysaccharide-stimulated succinate dehydrogenase-B degradation but not sequestration. We hypothesized that this Complex II regulatory mechanism might be coordinated by the mitochondrial phospholipid cardiolipin. Cardiolipin synthase knockdown prevented lipopolysaccharide-induced metabolic remodeling and Complex II disassembly, sequestration and degradation. Cardiolipin-depleted macrophages were defective in lipopolysaccharide-induced pro-inflammatory cytokine production, a phenotype partially rescued by Complex II inhibition. Thus, cardiolipin acts as a critical organizer of inflammatory metabolic remodeling.

## Introduction

Metabolic plasticity is a central feature of immunity (*1, 2*). Immune cells respond to diverse physiological cues, including infectious and sterile inflammatory stimuli, by eliciting distinct metabolic programs. While it has long been appreciated that these metabolic changes reshape cellular bioenergetics, more recent studies indicate that context-driven metabolic remodeling defines immune cell fate and function. Although metabolic plasticity is common among immune cells, the regulatory mechanisms and consequences of these metabolic changes are highly specialized in different cell types (*3*–*5*).

Metabolic remodeling in macrophages contributes to the initiation and resolution of inflammation in multiple human diseases, including sepsis, infection, inflammatory disease, and autoimmune disease (*3, 6*–*8*). Furthermore, a variety of metabolic determinants of inflammatory signaling have been identified in macrophages. Importantly, control of oxidative phosphorylation, particularly through modulation of respiratory chain (RC) function, is critical for inflammatory programming. Recent work has highlighted respiratory complex II (Complex II), which functions uniquely at the interface of the electron transport chain (ETC) and the tricarboxylic acid (TCA) cycle, as a major player in macrophage metabolic remodeling and inflammatory programming (*9*–*13*). While ample evidence connects Complex II and its substrate succinate to macrophage inflammatory programming, the regulatory mechanisms governing Complex II activity in macrophages remain to be defined.

Prior work points to accumulation and oxidation of succinate as critical for macrophage inflammatory programming. Inflammatory accumulation of succinate has been attributed to increased flux through the gamma-aminobutyric acid (GABA) shunt (*10*). Succinate produced through this mechanism enhances lipopolysaccharide (LPS)-stimulated inflammatory signaling through oxidative stress-dependent activation of the transcription factor Hypoxia Inducible Factor 1 (HIF-1), which increases the production of the immature form of the pro-inflammatory cytokine Interleukin 1β (pro-IL-1β). Nevertheless, sustained inhibition of Complex II, which results in succinate accumulation, suppresses inflammatory signaling. Notably, deletion of one of the four subunits of Complex II, succinate dehydrogenase B (SDHB), in macrophages leads to a hypo-inflammatory phenotype, as indicated by a defect in LPS-induced pro-IL-1β (*9*). Thus, current models support that Complex II is required for the inflammatory effect of succinate accumulation. Furthermore, acute bacterial infection of macrophages results in increased Complex II activity within the first two hours (*12*). Overall, the current literature predicts that precise and dynamic control of Complex II activity during inflammatory macrophage activation may be important, but early regulatory mechanisms controlling activity and stability of Complex II during the course of macrophage inflammatory programming are not well defined.

The supramolecular organization of the respiratory complexes (RCs) depends on phospholipids present in the inner mitochondrial membrane (IMM) (*14, 15*). In particular, the mitochondrial phospholipid cardiolipin (CL) scaffolds the RCs in supercomplexes and enhances their function. Furthermore, human diseases caused by defects in CL metabolism, such as Barth Syndrome, are associated with defects in RC organization and function across multiple cell types (*16*). While CL has been linked to RC function, little is known about how CL may control inflammatory modulation of RC activity in macrophages. Here, we identify regulation of SDHB as a key step in LPS-induced respiratory chain remodeling and demonstrate a critical role for CL in enabling Complex II disassembly and SDHB degradation. Our findings point to CL as an architect of macrophage metabolic remodeling that repurposes the respiratory chain towards inflammation.

## Results

### Macrophage inflammatory activation destabilizes Complex II subunit SDHB

Activation of Toll-like receptor 4 (TLR4) with lipopolysaccharide (LPS) triggers profound metabolic remodeling in macrophages which culminates in ATP production by aerobic glycolysis at the expense of oxidative phosphorylation (*3, 7, 13*). In agreement with prior studies, stimulation of murine macrophages with LPS led to decreased oxygen consumption (OCR) and increased extracellular acidification (ECAR) as early as 6h post-treatment, shown by Seahorse extracellular flux assay (**Fig. S1A-B**) (*6, 17, 18*). We therefore predicted that LPS stimulation might alter relative abundance or assembly of the respiratory complexes (RC). To test this hypothesis, we performed immunoblot analysis of representative RC subunits in macrophages stimulated with LPS. RC subunit analysis revealed selective loss of respiratory complex II (Complex II) subunit SDHB, but not SDHA, at 24h post-LPS stimulation (**Fig. 1A-B**). The other RCs remained at baseline levels or showed slight decreases that were not statistically significant. Our findings are consistent with earlier studies demonstrating that LPS stimulation triggers accumulation of the Complex II substrate succinate, complementing the production of succinate via the GABA shunt (*10*).

**Fig 1.**
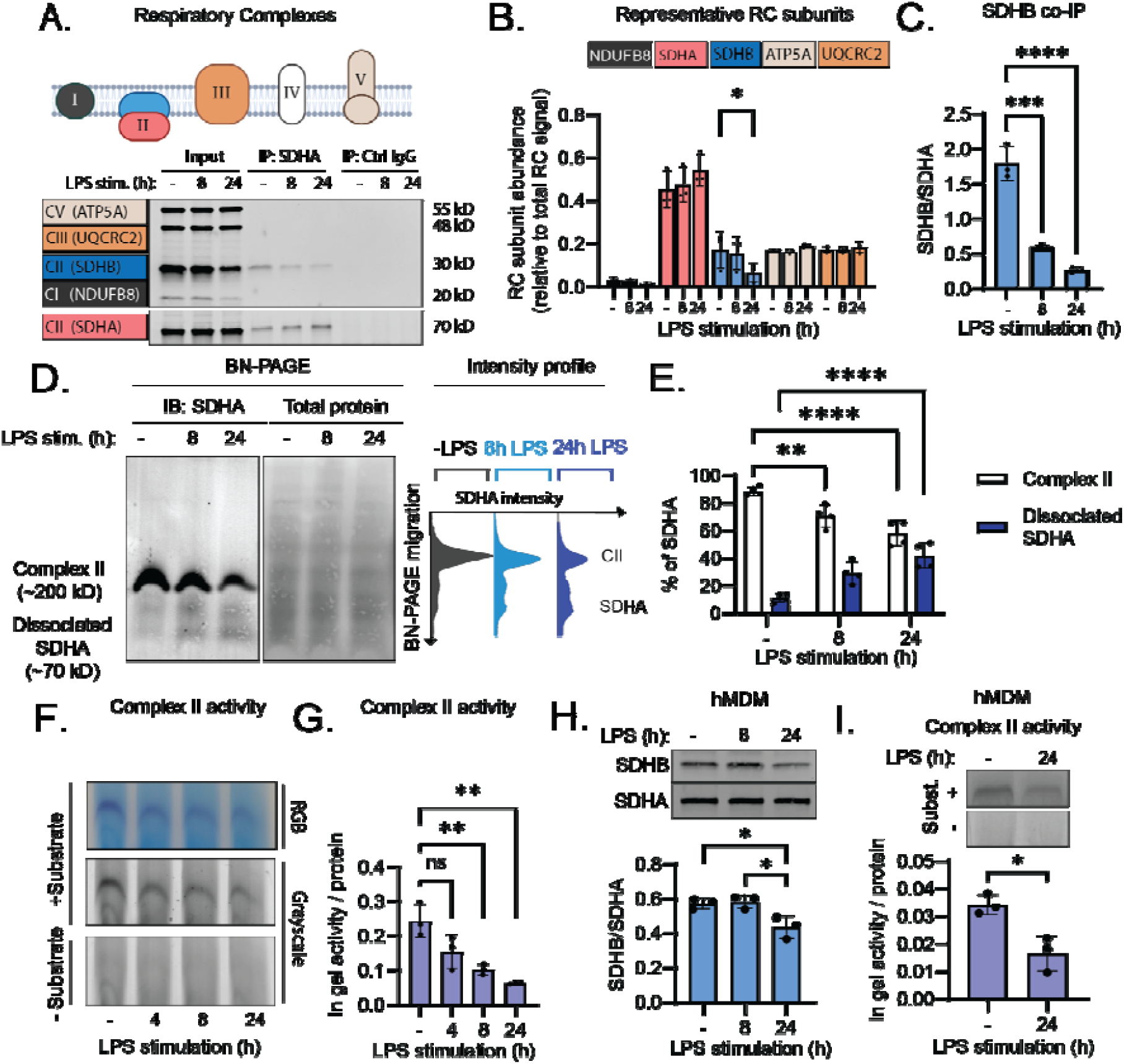
Macrophage LPS stimulation destabilizes Complex II. **A**. iBMDM were synchronously stimulated with or without 200 ng/mL LPS for 8 or 24h. Whole cell extracts were subject to SDHA or control IgG immunoprecipitation. Samples were analyzed by SDS-PAGE and immunoblot against a panel of respiratory complex (RC) subunits (ATPA5, UQCRC2, SDHA, SHDB, and NDUFB8). **B**. Quantification of RC subunit abundance from input sample from 1A relative to total RC signal. **C**. Quantification of immunoprecipitated SDHB relative to SDHA from 1A. **D**. Whole cell extracts from iBMDM synchronously stimulated with or without LPS for 8 or 24h were analyzed by blue native (BN) PAGE and subject to SDHA immunoblot. Intensity profile analysis of the SDHA blot shows two peaks, corresponding to Complex II-associated and dissociated SDHA. **E**. Quantification of the percentage of SDHA occupancy in Complex II and dissociated SDHA populations as highlighted in 1D. **F**. iBMDM were synchronously stimulated with or without LPS for 4, 8, or 24h and subject to a modified BN PAGE protocol (clear native [CN]-PAGE) and a succinate dehydrogenase in gel activity assay. **G**. Enzymatic activity quantified from each gel as in 1G was normalized to protein loading from the whole cell extract. **H**. Primary human monocyte-derived macrophages (hMDM) were stimulated with or without 200 ng/mL LPS for 8 or 24h. Whole cell extracts were subject to SDS-PAGE and SDHA/SDHB immunoblot. The ratio between SDHB and SDHA was quantified. **I**. Complex II succinate dehydrogenase activity was measured as in 1F-G from hMDM stimulated with or without LPS for 24h. Graphs are presented as the mean of n ≥ 3 independent experiments with standard deviation (SD) error bars. P values were calculated using an unpaired T-test or one-way ANOVA with Tukey’s post-test for multiple comparisons. *P < 0.05; **P < 0.01; ***P < 0.001.

To investigate how LPS treatment affects the assembly of Complex II, we immunoprecipitated SDHA and measured co-immunoprecipitation of other representative RC subunits (**Fig. 1A and C**). We found that, among the tested RC subunits, only SDHB co-immunoprecipitated with SDHA under unstimulated conditions, supporting a known phenomenon whereby Complex II rarely forms supercomplexes with other RCs (*19*). Importantly, LPS stimulation triggered dissociation of SDHA and SDHB as early as 8h post LPS stimulation. In support of this, blue native (BN)-PAGE analysis of Complex II during LPS stimulation showed a decrease in native Complex II, and a corresponding increase in dissociated SDHA, at 8 and 24h post LPS stimulation (**Fig. 1D-E)**. Notably, the assembly of other respiratory complexes was not perturbed by LPS stimulation (**Fig. S2**). Disassembly of Complex II correlated with decreased enzymatic activity, as indicated by an in-gel activity assay of Complex II succinate dehydrogenase activity (**Fig. 1F and G**). We next tested whether Complex II activity was regulated by a similar mechanism in primary human monocyte-derived macrophages (hMDM) and found that that indeed SDHB, but not SDHA, was decreased by LPS stimulation, resulting in loss of Complex II activity by 24h (**Fig. 1H and I)**. Finally, we found that pharmacological inhibition of Complex II was sufficient to trigger a glycolytic burst in macrophages (**Fig. S1C)**. Thus, we propose that Complex II disassembly and instability leads to functional inactivation, contributing to defective respiration and increased glycolysis during macrophage inflammatory activation.

### Complex II subunit SDHB is sequestered and selectively degraded through mitophagy during macrophage inflammatory activation

Selective loss of Complex II subunit SDHB, but not SDHA, upon LPS stimulation could be achieved by a variety of mechanisms, including gene expression and proteolytic pathways. To investigate SDHB protein-level regulation, we first explored the subcellular localization of Complex II during LPS stimulation. We visualized SDHA and SDHB subcellular localization by immunofluorescence staining and high resolution confocal fluorescence microscopy, counterstaining with Complex I subunit MT-ND1 (inner mitochondrial membrane) and LAMP1 (endolysosomal network) (**Fig. 2A and Fig. S3**). At 8h post-LPS stimulation, when we observed Complex II disassembly but not loss of SDHA or SDHB subunit abundance, we noted sequestration of SDHB into puncta that were distinct from the main mitochondrial network as defined by MT-ND1 immunofluorescence. In contrast, SDHA remained ubiquitous across the mitochondrial network (**Fig 2B-D, Fig S3B, Fig S4 C-D**). Interestingly, there was selective delivery of SDHB to the LAMP1-positive vesicular network (**Fig 2B and Fig S2A**). Optical sectioning and 3D rendering of confocal immunofluorescence micrographs revealed two subsets of SDHB puncta: one subset localized within spherical LAMP1 structures, while the other subset was retained within the mitochondrial network (**Fig S4A-B**). These results suggest that SDHB is sequestered within the mitochondrial network and then delivered to an endolysosomal compartment in response to LPS stimulation.

**Fig 2.**
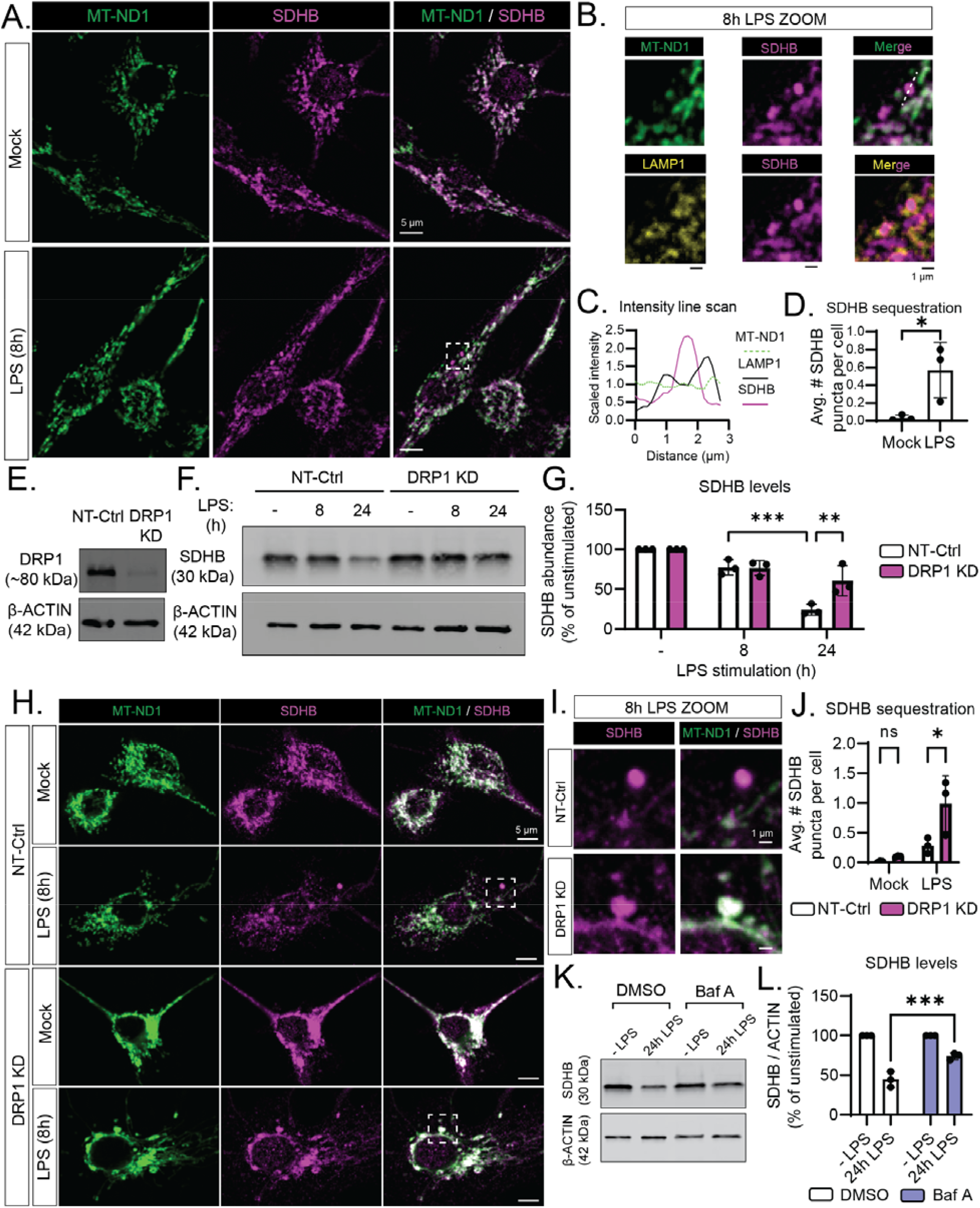
SDHB is degraded by selective mitophagy during LPS stimulation. Representative confocal fluorescence micrographs from WT iBMDM stimulated with or without 200 ng/mL LPS for 8h and subject to immunofluorescence labeling of Complex II subunit SDHB and Complex I subunit MT-ND1. **B**. Magnified region of interest from 2A and S3 highlighting sequestration of SDHB signal into puncta which are distinct from MT-ND1 signal and enclosed in a LAMP1-positive compartment. **C**. Representative line scan from SDHB puncta in 2B where pixel intensity is scaled to the mean intensity across the sampled region for each immunostain. **D**. Automated quantification of the average number of SDHB puncta per cell from 2A based on spot identification and thresholding modules in CellProfiler™. **E**. Whole cell extracts from iBMDM expressing non target control (NT-Ctrl) or Dynamin-Related Protein 1 (*Drp1*)-targeted shRNA (DRP1 KD) were subject to SDS-PAGE and immunoblot analysis of DRP1. **F**. NT-Ctrl and DRP1 KD iBMDM were stimulated with or without 200 ng/mL LPS for 8 or 24h and SDHB levels were assessed by SDS-PAGE and immunoblot. **G**. Quantification of 2F as percentage of the unstimulated condition (-LPS) within each cell line. **H**. Representative confocal fluorescence micrographs from NT-Ctrl and DRP1 KD iBMDM stimulated with or without 200 ng/mL LPS for 8h and subject to immunofluorescence labeling of SDHB and MT-ND1. **I**. Magnified region of interest from 2H highlighting retention of LPS-induced SDHB puncta within MT-ND1 signal in DRP1 KD iBMDM. **J**. Automated quantification of the average number of SDHB puncta per cell from 2H, as described in 2D. **K**. iBMDM were treated with or without 200 ng/mL LPS with either Bafilomycin A1 (Baf A, 100 μM) or vehicle control (0.1% DMSO). SDHB levels from whole cell extracts were measured by SDS-PAGE and immunoblot. **L**. Quantification of 2K as percentage of the unstimulated condition (-LPS) within each treatment condition. Graphs are presented as the mean of n = 3 independent experiments with standard deviation (SD) error bars. For image analysis, the mean of ∼100 cells per condition is reported for each independent experiment. P values were calculated using an unpaired T-test or two-way ANOVA with Sidak’s post-test for multiple comparisons. *P < 0.05; **P < 0.01; ***P < 0.001.

Delivery of mitochondrial components to the endolysosomal network can occur during mitochondrial autophagy (mitophagy), so we reasoned that the decrease in SDHB protein abundance might be mediated by mitophagy. During mitophagy, dynamin-related protein (DRP1)-dependent mitochondrial fission generates small (∼1 μm^2^), fragmented mitochondria which can be recognized by the autophagy machinery and captured for disposal. Thus, we predicted that DRP1 depletion would inhibit LPS-induced SDHB delivery to a LAMP1-positive compartment. We therefore stimulated DRP1 KD macrophages (**Fig. 2E**) (*20*) with LPS, and measured SDHB levels by immunoblot **(Fig. 2F-G**). LPS-stimulated DRP1 KD macrophages failed to exhibit decreased SDHB compared to cells expressing non-target control shRNA (NT-Control). Additionally, we tracked SDHB subcellular localization by immunofluorescence labeling and confocal microscopy (**Fig. 2H**). We observed more prominent SDHB puncta formation in LPS-stimulated DRP1 KD macrophages, but these puncta remained associated with the mitochondrial network (**Fig. 2I-J**) Thus, we find that DRP1 KD macrophages are capable of sequestering SDHB but infer that they have a defect in their ability to release SDHB puncta from the mitochondrial network for degradation. To test whether endolysosomal degradation could contribute to LPS-induced loss of SDHB, we treated cells with LPS in the absence or presence of the vacuolar ATPase inhibitor Bafilomycin A1 (Baf A) and measured SDHB levels by immunoblot (**Fig 2K-L**). We found that SDHB was not as efficiently lost in response to LPS when Baf A was present, compared to vehicle control. While our data support selective mitophagy of SDHB, we also tested whether differences in *Sdhb* transcript could account for differences in protein levels. We stimulated macrophages with LPS and measured expression of *Sdha, Sdhb, Sdhc*, and *Sdhd* by RT-qPCR (**Fig. S5**). We found that LPS stimulation led to a slight (trending) decrease in transcript levels of all Complex II subunits except *Sdhb*, indicating that this mechanism is unlikely to account for selective loss of SDHB. Taken together, our data support Complex II disassembly, followed by SDHB sequestration and degradation, as a functional consequence of LPS-induced macrophage innate immune signaling.

### CL licenses metabolic remodeling during macrophage inflammatory activation

As SDHB appears to be selectively degraded through mitophagy when macrophages are stimulated with LPS, we sought to determine the mechanism by which SDHB is selectively packaged and degraded while other mitochondrial proteins, like SDHA, remain unperturbed. Prior work has determined that the mitochondrial phospholipid cardiolipin (CL) regulates mitophagy upon exposure to the mitochondrial surface, a phenomenon known to occur during LPS stimulation (*21, 22*). Furthermore, CL facilitates supramolecular organization and function of the respiratory complexes (*23*). Thus, we hypothesized that CL proximity to and/or interaction with Complex II might control selective mitophagy of SDHB during LPS stimulation. To investigate the role of CL in macrophage metabolism, we knocked down the terminal enzyme in the CL biosynthetic pathway, Cardiolipin Synthase (CRLS1 KD), and validated an effect on CRLS1 and CL levels by immunoblot and untargeted lipidomics of mitochondrial fractions, respectively (**Fig. 3A-B; Fig S6**). We noted that CRLS1 KD did not significantly affect the basal abundance of respiratory complex subunits, respiration, glycolysis, or membrane potential (**Fig 3C-F; Fig S7)**. These findings suggest that the minimum amount of CL needed for basal metabolism is achieved in CRLS1 KD macrophages. While CRLS1 KD macrophages maintained apparently normal basal metabolism, disruption of CL biosynthesis substantially interrupted the ability of macrophages to dampen respiration and switch to aerobic glycolysis during LPS stimulation (**Fig. 3D-F**). Consistent with a failure to inhibit respiration, CRLS1 KD macrophages did not increase mitochondrial reactive oxygen species (mtROS) during LPS stimulation, evidenced by decreased intensity of the mitochondrial superoxide indicator, MitoSOX, quantified by confocal imaging (**Fig. 3G-H**). Collectively, these data suggest that the ability of macrophages to remodel metabolism in response to LPS stimulation depends on CL biosynthesis.

**Fig 3.**
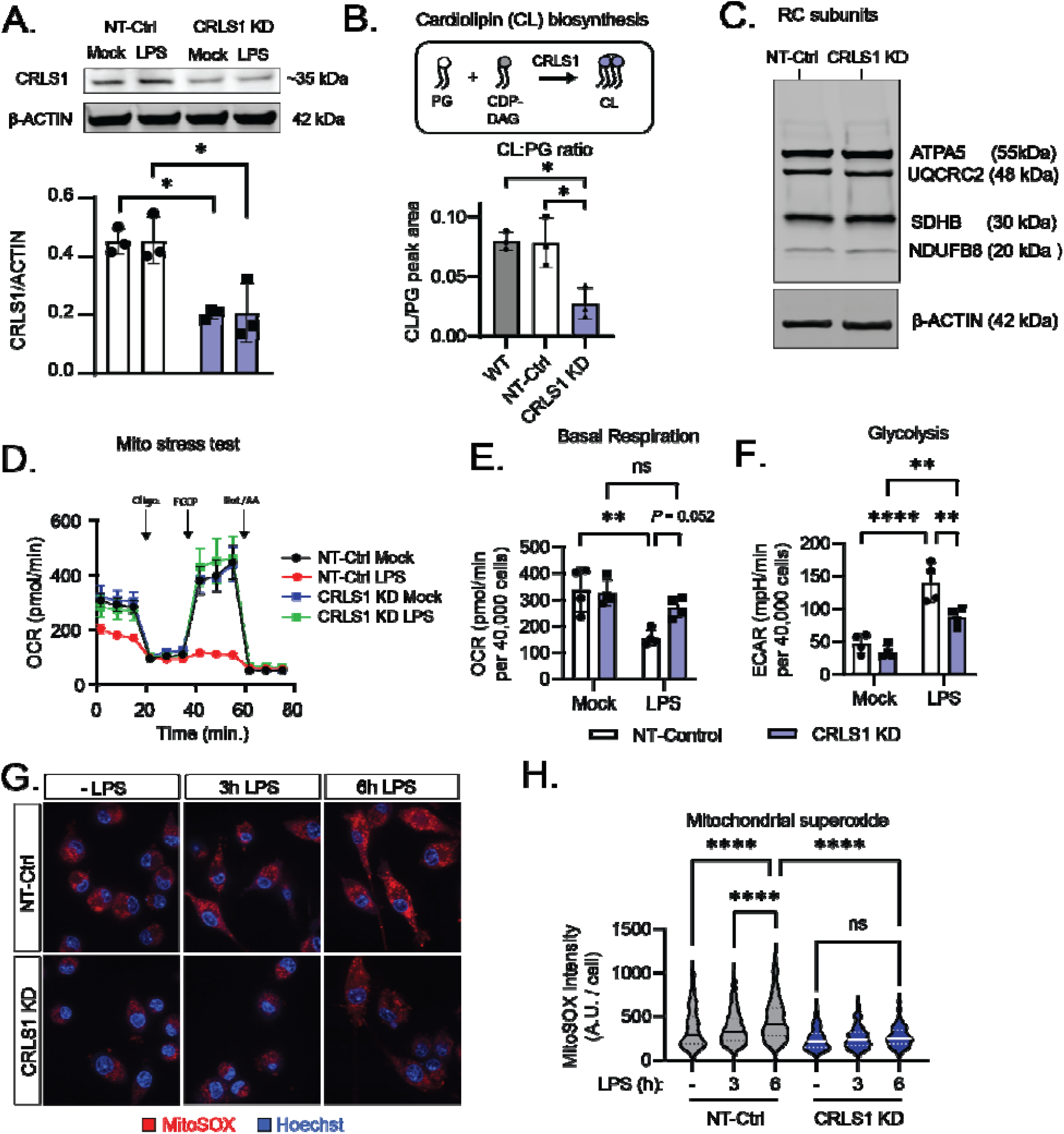
CL biosynthesis licenses metabolic remodeling during macrophage LPS stimulation. **A**. Whole cell extracts from iBMDM expressing non-target control (NT-Ctrl) or Cardiolipin Synthase (*Crls1*)-targeted shRNA (CRLS1 KD) stimulated with or without 200 ng/mL LPS for 6h were subject to SDS-PAGE and immunoblot analysis of CRLS1. **B**. Cartoon showing cardiolipin synthesis from phosphatidylglycerol (PG) and cytidine diphosphate diacylglycerol (CDP-DAG) in the mitochondria. Untargeted lipidomics was performed on WT, NT-Ctrl, and CRLS1 KD iBMDM mitochondrial isolates. Ratio of peak area corresponding to total measured species of cardiolipin (CL) and its precursor phosphatidylglycerol (PG) is reported. **C**. NT-Ctrl and CRLS1 KD iBMDM were subject SDS-PAGE and immunoblot analysis of a panel of representative RC subunits (ATPA5, UQCRC2, SDHA, SDHB, and NDUFB8). **D**. NT-Ctrl and CRLS1 KD iBMDM were pretreated with 200 ng/mL LPS for 6h and then subject to the Agilent Seahorse Extracellular Flux (XF) analysis of the oxygen consumption rate (OCR) and the extracellular acidification rate (ECAR) during the Mito Stress Test assay. The following concentrations of respiratory chain inhibitors were used in the Mito Stress Test: 2 μM FCCP, 1.5 μM oligomycin, 0.5 μM rotenone, and 0.5 μM antimycin A. **E**. Quantification of basal respiration (OCR) as in 3D across 4 independent experiments. **F**. Quantification of basal ECAR as a readout of glycolysis across 4 independent experiments corresponding to 3D and 3E. **G**. Representative confocal fluorescence micrographs from NT-Ctrl and CRLS1 KD iBMDM synchronously stimulated with or without 200 ng/mL LPS for 3 or 6h and then stained with the mitochondrial superoxide indicator MitoSOX and counterstain Hoechst. **H**. Automated quantification of MitoSOX intensity per cell from μ300 cells across 3 independent experiments, as shown in 3G, using Cell Profiler™. Graphs are presented as the mean of n ≥ 3 independent experiments with standard deviation (SD) error bars. For image analysis, the mean of ∼100 cells per condition is reported for each independent experiment. P values were calculated using a one-way ANOVA with Tukey’s post-test for multiple comparisons or two-way ANOVA with Sidak’s post-test for multiple comparisons. *P < 0.05; **P < 0.01; ***P < 0.001; ****P < 0.0001.

### CL biosynthesis is required for Complex II destabilization and sequestration during macrophage inflammatory activation

Macrophage stimulation with LPS triggers rewiring of the TCA cycle, a process which is linked to inflammatory macrophage polarization (*24*). Since we identified a key role of CL biosynthesis in global metabolic remodeling in LPS-stimulated macrophages, we tested whether CRLS1 KD macrophages were defective in their capacity to alter TCA cycle metabolites in response to LPS. To this end, we performed a metabolomics kinetic study to identify changes in metabolite levels between CRLS1 KD and NT-control macrophages during LPS stimulation. Our analysis revealed that LPS triggers a CRLS1-dependent break in the TCA cycle, whereby succinate accumulates and downstream metabolites malate and aspartate are depleted (**Fig 4A and Fig S8**). Additionally, we observed decreased accumulation of itaconate in LPS-stimulated CRLS1 KD macrophages, a phenotype normally associated with Complex II dysfunction (**Fig S8**) (*25*). As succinate is a Complex II substrate, we hypothesized that CRLS1 KD macrophages might have a defect in Complex II stability or activity. We first tested if CL contributes to LPS-induced destabilization of Complex II subunit SDHB. To this end, we stimulated macrophages with or without LPS for 24h and measured abundance and activity of Complex II by BN-PAGE and parallel immunoblot and in-gel activity assays (**Fig. 4B-E**). CL biosynthesis was critical for LPS-induced disassembly and inhibition of Complex II. Furthermore, SDS-PAGE and immunoblot analysis of RC subunits revealed that SDHB was lost in NT-Control macrophages, but retained in CRLS1 KD macrophages (**Fig. 4F-G**). Since Complex II disassembly was prevented in CRLS1 KD macrophages, we predicted that SDHB sequestration would also be limited. To test this hypothesis, we tracked localization of MT-ND1 and SDHB by immunofluorescence and high resolution confocal microscopy. We found that CL biosynthesis was required for focal accumulation of SDHB in response to LPS stimulation, since CRLS1 KD macrophages failed to produce SDHB puncta (**Fig. 4H-I**). Together, our data support that LPS-induced SDHB sequestration and degradation through mitophagy depend on CL.

**Fig 4.**
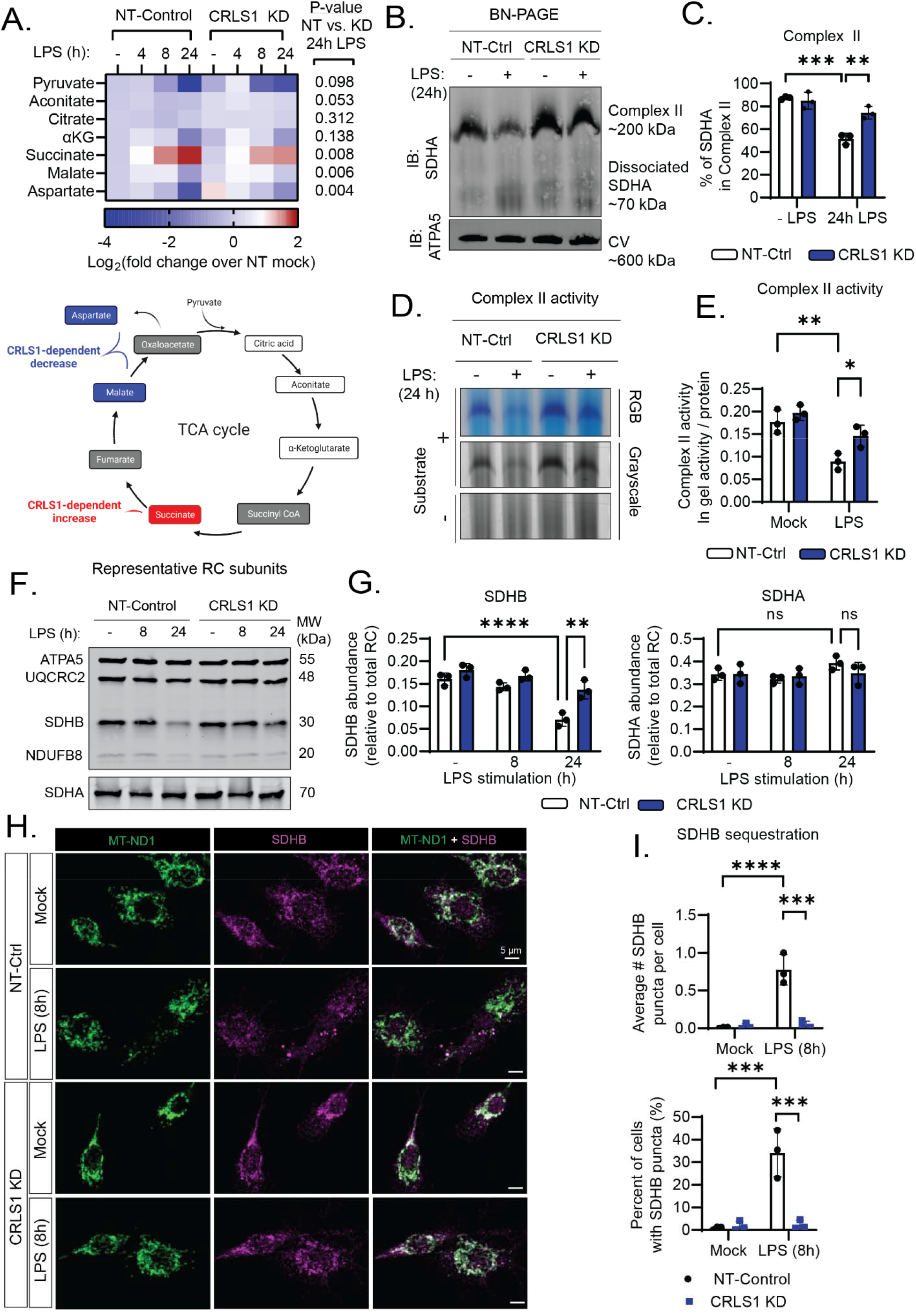
CL biosynthesis is required for Complex II disassembly and degradation during LPS stimulation. **A**. Highlighted TCA cycle metabolites from targeted metabolomics analysis of NT-Ctrl and CRLS1 KD iBMDM synchronously stimulated with or without 200 ng/mL LPS for 4, 8, or 24h. **B**. Whole cell extracts from NT-Ctrl and CRLS1 KD iBMDM stimulated with or without LPS for 24h were analyzed by blue native (BN) PAGE and subject to SDHA and ATPA5 immunoblot. **C**. Quantification of the percentage of SDHA occupancy in Complex II from 4B, as performed in 1E. **D**. CN-PAGE and in gel succinate dehydrogenase activity assay of NT-Ctrl and CRLS1 KD iBMDM stimulated with or without LPS for 24h. **E**. Quantification of 4D as in gel activity normalized to protein loading. **F**. SDS-PAGE and immunoblot analysis of representative RC subunits of NT-Ctrl and CRLS1 KD iBMDM synchronously stimulated with 200 ng/mL LPS for 8 or 24h. **G**. Quantification of SDHB and SDHA abundance relative to total RC signal from 4F. **H**. Representative confocal fluorescence micrographs from NT-Ctrl and CRLS1 KD iBMDM stimulated with or without 200 ng/mL LPS for 8h and subject to immunofluorescence labeling of SDHB and MT-ND1. **I**. Automated quantification of the average number of SDHB puncta per cell and percentage of cells with SDHB puncta from 4H, as described in 2D. Graphs are presented as the mean of n = 3 independent experiments with standard deviation (SD) error bars. For image analysis, the mean of ∼100 cells per condition is reported for each independent experiment. P values were calculated using a two-way ANOVA with Sidak’s post-test for multiple comparisons. *P < 0.05; **P < 0.01; ***P < 0.001; ****P < 0.0001.

### CL biosynthesis and modulation of Complex II activity are critical for early inflammatory responses in macrophages

Complex II and CL have been implicated in macrophage inflammatory function through HIF-1-driven inflammatory gene expression (*10, 22*). With this observation in mind, we hypothesized that interaction between CL and Complex II may regulate macrophage inflammatory programming. To test our hypothesis, we stimulated NT-Control and CRLS1 KD with LPS and measured transcript levels of hallmark inflammatory genes *Il6* and *Tnf* by RT-qPCR. CRLS1 KD macrophages exhibited markedly decreased levels of *Il6* transcript compared to NT-control, with only a trending defect in transcript levels of *Tnf* (**Fig 5A**). In parallel, we measured secreted levels of these cytokines by ELISA (**Fig 5B**). Similar to transcript levels, we found that CRLS1 was required for LPS-induced production of IL-6, but not TNF-α. To determine if CRLS1 was important for inflammatory responses in other types of macrophages beyond bone marrow-derived macrophages, we knocked down CRLS1 in the peritoneal origin RAW264.7 murine macrophage cell line and saw similar CRLS1-dependent inflammatory responses (**Fig. S9**). Finally, we observed that CRLS1 KD in macrophages prevented IL-6 production in response to the bacterial pathogen *Salmonella enterica* Typhimurium (**Fig S10)**. These findings indicate that CL biosynthesis selectively regulates inflammatory gene expression at the transcript level.

**Fig 5.**
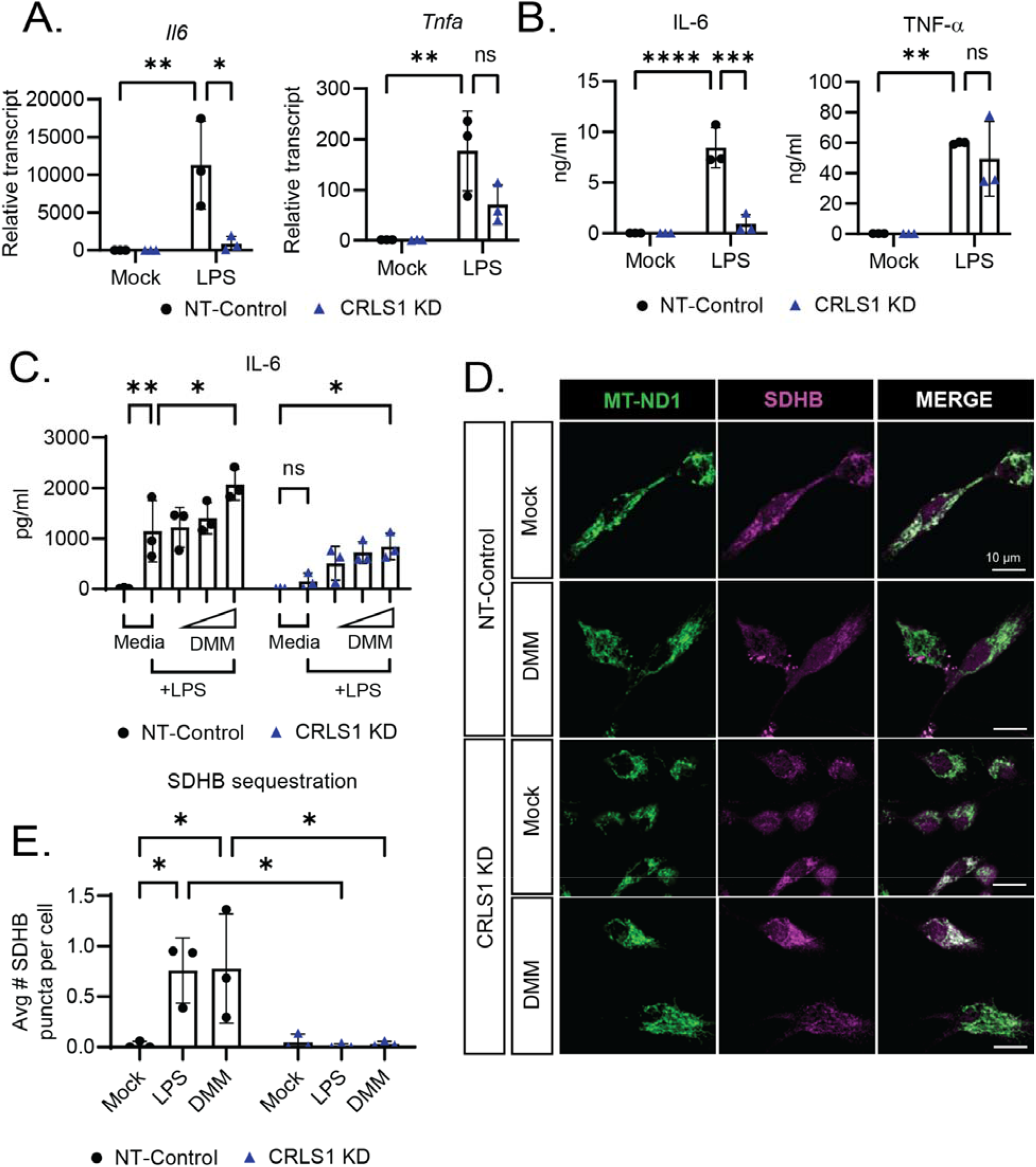
CL biosynthesis and Complex II inhibition contribute to inflammatory gene expression during LPS stimulation. **A**. RT-qPCR analysis of transcript levels of *Il6* and *Tnf* in NT-Ctrl and CRLS1 KD iBMDM stimulated with or without 200 ng/mL LPS for 4h. **B**. ELISA analysis of IL-6 and TNF-α secreted by NT-Ctrl and CRLS1 KD iBMDM stimulated with 200 ng/mL LPS for 24h. **C**. ELISA analysis of IL-6 secreted by NT-Ctrl and CRLS1 KD iBMDM pretreated for 1h with or without 0.1, 1, or 10 mM dimethyl malonate (DMM) then challenged with 200 ng/mL LPS for 6h. **D**. Representative confocal fluorescence micrographs from NT-Ctrl and CRLS1 KD iBMDM stimulated with or without 10 mM DMM for 8h and subject to immunofluorescence labeling of SDHB and MT-ND1. **E**. Quantification of SDHB puncta in NT-Ctrl and CRLS1 KD iBMDM stimulated with 10 mM DMM or 200 ng/mL LPS for 8h, quantified as described in Fig 2D. Complete set of representative confocal fluorescence micrographs shown in Fig S12. Graphs are presented as the mean of n = 3 independent experiments with standard deviation (SD) error bars. For image analysis, the mean of ∼100 cells per condition is reported for each independent experiment. P values were calculated using a two-way ANOVA with Sidak’s post-test for multiple comparisons. *P < 0.05; **P < 0.01; ***P < 0.001; ****P < 0.0001.

Our results provide evidence that CL biosynthesis is critical for inflammatory programming in macrophages, and work from other labs collectively supports that Complex II regulates inflammatory responses in macrophages. To test if these are related processes, we treated LPS-stimulated NT-control or CRLS KD macrophages with two different commercially available Complex II inhibitors, Atpenin A5 (AA5) or dimethyl malonate (DMM), at a range of sub-cytotoxic concentrations and measured IL-6 production by ELISA (**Fig 5C and Fig S11**). We found that both of these Complex II inhibitors partially restored IL-6 production in CRLS1 KD macrophages. We conclude that CL biosynthesis is critically important for inflammatory programming in macrophages, and that there are likely Complex II-dependent and -independent components to this programming. Interestingly, exogenous addition of diethyl succinate did not rescue IL-6 production, indicating that modulation of Complex II activity may be more nuanced than accumulation of succinate for inflammatory outcomes in the early phase of inflammatory activation. (**Fig S12**). Finally, we tested whether inhibition of Complex II was sufficient to trigger SDHB sequestration. We found that DMM-treated macrophages sequestered SDHB into puncta to a similar extent as LPS-treated cells (**Fig 5D-E and Fig S13**). Additionally, DMM-induced SDHB sequestration was dependent on CRLS1. Thus, we speculate that CL biosynthesis contributes to separable processes of Complex II disassembly and sequestration of dysfunctional Complex II subunits.

## Discussion

TLR4 activation by LPS triggers global metabolic remodeling, shifting away from respiration and toward glycolysis in macrophages, repurposing the respiratory chain for host defense and inflammatory signaling. Inhibition of the respiratory complexes themselves triggers substantive metabolic reprogramming and is sufficient to stimulate a robust glycolytic burst (*26*). From these observations, we predicted that one or more of the RCs would be negatively regulated by LPS stimulation. We found that Complex II activity decreased and its components disassembled in response to LPS stimulation. Following disassembly, Complex II subunit SDHB was sequestered into microdomains of the mitochondrial network and released in a DRP1-dependent manner, reminiscent of mitophagy. SDHB released from the mitochondrial network through this mechanism was turned over through endolysosomal degradation. The mitochondrial phospholipid cardiolipin (CL) was critical for disassembly of Complex II, and sequestration and degradation of SDHB. Finally, we show that defective CL biosynthesis compromises the induction or stability of pro-inflammatory cytokine transcripts, notably IL-6, a phenotype which could be partially rescued by Complex II inhibition. Collectively, our work highlights a novel function for CL in regulating macrophage inflammatory programming through coordination of Complex II disassembly and degradation.

Ample evidence supports an integral role for regulation of electron transport chain components in executing the shift from homeostatic to inflammatory programming (*27*). Environmental cues alter macrophage metabolism which guides polarization between inflammatory and anti-inflammatory macrophage phenotypes. Metabolic changes influence cellular fate through a variety of factors, including oxidative stress pathways, post-translational modification of proteins by induced metabolites, and epigenetics. O’Neill and colleagues demonstrated that succinate produced through the GABA shunt enhanced LPS-induced IL-1β production, while conditional deletion of the *Sdhb* gene in macrophages or pharmacological inhibition of Complex II, both approaches that increase succinate pools, paradoxically compromised IL-1β production during prolonged LPS stimulation (*9*). The model that best fits these data is that succinate oxidation by Complex II participates in inflammatory signaling. Nevertheless, our data demonstrate that pharmacological inhibition of Complex II in the acute early phase of LPS stimulation enhances inflammatory responses, particularly IL-6 production. Furthermore, our data demonstrate that succinate oxidation alone cannot account for the inflammatory contribution of Complex II, as pro-inflammatory cytokine production in CRLS1 KD macrophages, which maintain succinate dehydrogenase activity, cannot be rescued by exogenous succinate supplementation. Together, our work and prior studies point to dynamic regulation of both Complex II and succinate levels as critical for orchestrating both the acute inflammatory response as well as the subsequent shift from pro- to anti-inflammatory macrophage function. Indeed, prior work has identified that alternative assembly of Complex II, reminiscent of what we have observed, aids in the adaptation to energetic stress (*28*). Following the model of alternative assembly, we propose that the SDHA/SDHB dimer dissociates from the integral membrane SDHC/SDHD dimer to alter Complex II function. In fact, at 8h post LPS stimulation, we observed by BN-PAGE and SDHA immunoblot a banding pattern consistent with an SDHA/SDHB dimer (**Fig S2**). Subsequently, by 24h post LPS stimulation we have observed loss of SDHB but not SDHA, indicating further dissociation of the SDHA/SDHB dimer and selective loss of SDHB. Thus, we propose that later loss of SDHB may be a mechanism to dampen the inflammatory function of the SDHA/SDHB dimer, though further studies are needed to test these hypotheses.

Recent findings indicate that inhibitors of Complex I, II, III or V prevent NLRP3 inflammasome activation and IL-1β production, by sustaining high ATP levels in a ROS-independent manner (*29*). Although observations using pharmacological inhibition must be interpreted with caution, the preponderance of data support that components of the respiratory complexes are prime targets for multi-level regulation. In addition to alterations in transcript or protein levels, Clayton et al, determined that in response to inflammatory signals, the Cytochrome *c* oxidase (C*c*O) subunit NDUFA4, is replaced by a paralogous component, C15ORF48 (*30*). The expression of *C15orf48* transcript is positively correlated with disease severity in patients with rheumatoid arthritis. In addition, macrophages from human patients genetically lacking NDUFA4, a subunit of CIV, exhibit hyperinflammatory characteristics, independent from changes in cellular ATP production rates, again underscoring the complex relationship between the function of respiration and its component parts (*30*). Our findings identify a regulatory modality that controls availability and function of SDHB protein during the early pro-inflammatory macrophage response, thereby tuning the activity of CII. Our results further establish that cardiolipin, a key architect of homeostatic mitochondrial function, plays a prominent and distinct role in CII regulation, and possibly other respiratory complexes, during LPS-induced mitochondrial reprogramming that leads to inflammation.

Metabolic reprogramming is implicated in the pathophysiology of many human diseases, with changes in respiratory chain function evident in many disease states. Genes encoding components of Complex II are mutated in several types of cancer, including hereditary paraganglioma due at least in part to increased ROS production and cell proliferation (*31*–*33*). In β-cells, deficiency in Complex II leads to metabolic dysfunction and diabetes in a mouse model of disease (*34*). A substantial body of evidence therefore points to Complex II as a critical regulatory nexus for metabolic reprogramming, and accordingly positions it as a target of multiple layers of regulation. Most work to date has focused on transcriptional regulation and post-translational modifications as regulatory determinants of Complex II abundance and activity. Post-translational regulatory mechanisms, particularly degradative pathways, have been identified for other respiratory complexes, including multiple subunits of Complex I, but not Complex II(*35*). Finally, the activity, abundance, and assembly of RCs are affected under diverse biological contexts. Thus, our observations that Complex II stability and activity is disrupted by TLR4 activation in macrophages fits into the broader context of complex regulation of the RCs. We propose that assembly-level control of Complex II enables rapid modulation of inflammatory metabolism in macrophages. Notably, we observe a trending drop in Complex II activity as early as 4h post LPS stimulation and a substantial drop in respiration by 6h post LPS treatment. Further, these early metabolic changes may regulate gene expression during inflammatory signaling. In fact, LPS-induced TCA cycle remodeling has been shown to alter inflammatory signaling through post-translational modifications of signaling molecules and epigenetic decoration of histones by TCA intermediates (*36*). The multi-protein respiratory complexes present many molecular interfaces for post-translational regulation and how such mechanisms determine the nature and magnitude of inflammation is ripe for further study.

Different inflammatory stimuli elicit unique metabolic and inflammatory programs in macrophages (*3*). These metabolic changes are regulated by complex and multifactorial mechanisms(*37*). One particular mechanism for the regulation of mitochondrial enzymes is control of protein abundance. The abundance of mitochondrial proteins can be regulated by a variety of mechanisms including gene expression, efficiency of mitochondrial import, degradation by local proteases, and organelle-level autophagic turnover of mitochondria (mitophagy). Here, we provide evidence that degradation of the Complex II subunit SDHB occurs through a novel route of selective mitophagy, whereby specific cargoes are enriched into fragmented mitochondria and targeted for endolysosomal degradation. Prior work from our lab and others has identified that TLR4 activation triggers the generation of a pool of small, fragmented mitochondria through the activity of Dynamin-related Protein 1 (DRP1)(*20, 38*). While the purpose of these fragmented mitochondria was previously unclear, data from this study supports that a subset of these fragmented mitochondria are enriched with specific mitochondrial cargo, including SDHB, to be turned over through mitophagy. CL organizes the mitochondrial inner membrane into functional microdomains, where it can facilitate respiratory supercomplex assembly and function(*14*). Our CRLS1 KD macrophages did not have a respiratory defect. Remarkably, one CL species was preferentially preserved even in the context of CRLS1 KD (**Fig. S6**). This particular acyl chain state is maintained by a CL-specific acyltransferase Tafazzin (TAZ)(*39*). Thus, we speculate that preservation of this CL species is sufficient to sustain homeostatic respiratory chain function. Alternatively, normal respiration in the CRLS1 KD macrophages may be attributed to an increase in phosphatidylethanolamine (PE). PE has been shown to support RC function and a compensatory increase in this phospholipid may sustain RC function when CL is limiting(*15*).

CRLS1 deficiency did not disrupt homeostatic respiration in macrophages, but these cells failed to remodel their metabolism in response to LPS stimulation, pointing to a key role for CL specifically in metabolic plasticity. Importantly, we observed that CL biosynthesis is required for the glycolytic burst and the production of mitochondrial superoxide, both of which are linked to inflammatory signaling in macrophages(*9, 10*). Furthermore, we found that CL biosynthesis was required for selective inflammatory gene expression, where LPS-induced *Il6* transcript levels were more sensitive to CL biosynthesis than *Tnf* transcript levels. This effect was partially dependent on CL-dependent modulation of Complex II stability, but was not completely restored by pharmacological Complex II inhibition. Our results indicate that CL biosynthesis contributes to inflammatory signaling through multiple routes. In fact, CL which is exposed to the OMM through membrane contact sites generated by NME4 contributes to inflammatory signaling upstream of NF-kB(*40*). Thus, CL may contribute to metabolic changes and inflammatory signaling cascades to strictly control inflammatory responses at the transcript level. Additionally, CL is proposed to scaffold the assembly of the NLRP3 inflammasome (*22*). Thus, CL contributes to inflammatory programming in macrophages at multiple levels. Within this context, we have identified a fundamental role for CL in early metabolic changes in macrophages which likely shapes later processes of gene expression and post-translational regulation of inflammatory responses. Taken together, our work reveals a novel mechanism by which macrophages adapt their metabolism to inflammatory cues. We have identified a CL-dependent route by which Complex II is disassembled and destabilized in macrophages. Further, our work has revealed that CL enables selective mitophagy of SDHB during inflammatory macrophage activation, without depleting its partner, SDHA. We speculate that CL microdomains may more generally provide the spatial architecture for selective packaging of mitochondrial cargo, acting as a key platform to execute different metabolic programs.

## Materials and methods

### Ethics statement

All animals used for experimental protocols were housed in specific pathogen free facilities at the University of Michigan Medical School Unit for Laboratory Animal Medicine (ULAM) and treated humanely in accordance with an IACUC-approved protocol. Blood samples were obtained from healthy adult donors according to the protocol approved by the University of Michigan Medical School (HUM00044257). Written consent was obtained from all donors.

### Cell culture

Murine immortalized bone marrow-derived macrophages (iBMDMs) were generated as previously described (*41*–*43*). Briefly, recombinant Cre-J2 virus containing v-raf and v-myc oncogenes was generated in 3T3 fibroblasts grown in Dulbecco’s Modified Eagle Medium (DMEM) supplemented with 10% heat-inactivated fetal bovine serum (FBS) and 50 U/ml of Penicillin and 50 μg/ml of Streptomycin. Sterile-filtered culture supernatants containing Cre-J2 virus were stored at −80°C. C57BL/6J lineage mouse femurs and tibiae were flushed and cells were transduced with Cre-J2 virus in macrophage differentiation media (50% DMEM, 2 mM L-Glutamine, 1 mM Sodium Pyruvate, 30% L929 cell-conditioned medium, 20% FBS, 50 U/ml of Penicillin and 50 μg/ml of Streptomycin). iBMDM were grown for at least 1 month before use in experiments to ensure immortalization was successful. L-929 cells were cultured in Minimum Essential Eagle Medium (MEM) supplemented with 2 mM L-glutamine, 1 mM sodium pyruvate, 1 mM non-essential amino acid (NEAA), 10 mM HEPES, and 10% FBS. All experiments were performed in DMEM supplemented with 2 mM L-glutamine, 1 mM Sodium Pyruvate and 10% FBS, unless otherwise indicated. When indicated, macrophages were treated with 200 ng/mL lipopolysaccharide (LPS) derived from *Salmonella enterica* serovar Typhimurium (Sigma-Aldrich, L2262). Additionally, a variety of inhibitors were used in this study, the concentration and source of which is included in the appended table. Throughout culturing, all cells were incubated at 37°C in 5% CO2.

### Human macrophage differentiation

Blood samples were obtained from healthy adult donors. Peripheral blood mononuclear cells were isolated by ficoll separation and differentiated *in vitro* into human monocyte-derived macrophages (hMDM). Briefly, blood was layered on ficoll and centrifuged at 500 x g at RT with low acceleration and no breaks for 20 min. Plasma was discarded, and the buffy coat and the layer beneath were collected, centrifuged, and resuspended in RPMI 1640 containing 20% FBS and 50 ng/ml human macrophage colony-stimulating factor (M-CSF) (300-25, Peprotech). Cells were seeded into sterile tissue-culture plates and differentiated for 7 days. At 3 days post-isolation, cultures were supplemented with fresh media. On the day of the experiment, cells were washed 3 times with PBS without Ca^2+^ or Mg^2+^ to remove non-adherent cells. Differentiation of macrophages by this protocol was validated by detection of CD68 and CD11b double-positivity by flow cytometry.

### Bacterial infection

*Salmonella enterica* serovar Typhimurium (STM) (**SL1344**) was grown overnight in LB in a 37°C slanted, shaking incubator (250 rpm). Bacteria were washed three times and diluted in sterile Dulbecco’s phosphate buffered saline (DPBS). iBMDM were infected with STM at an MOI of 20, and extracellular bacteria were killed with high concentration gentamicin treatment (100 μg/ml) for 1h, washed with PBS, and then low concentration gentamicin (10 μg/ml) was maintained for the remaining 23h of infection.

### Generation of DRP1 KD and CRLS1 KD

Stable knock down (KD) of Dynamin-related Protein 1 (DRP1) and Cardiolipin Synthase (CRLS1) in iBMDM was achieved using lentiviral delivery of shRNA. Lentivirus was generated and packaged in HEK293T cells grown in DMEM supplemented with 10% FBS. HEK293T cells were transfected with pLKO.1 plasmid encoding *Drp1*-targeted shRNA, *Crls1*-targeted shRNA, and a non-target control (NT-Ctrl) along with the packaging plasmids (pHCMV-G, and pHCMV-HIV-1) (*44*) using FUGENE-HD transfection reagent (Promega). The mouse *Drp1*-targeted shRNA plasmid with the sense sequence of GGCAATTGAGCTAGCTATA, *Crls1*-targeted shrRNA plasmid with the sense sequence of GAAGACTTTAATGTTGCACTA, and the non-target control shRNA plasmid were purchased from Sigma-Aldrich. Lentivirus-containing supernatants were collected, filtered, and used to transduce macrophage cell lines. Transduced cells were selected with puromycin (3 μg/ml) and resistant cells were grown and used for the experiments.

### Mitochondrial isolation

Mitochondria were isolated from cells by subcellular fractionation and density centrifugation as previously described(*45*). Briefly, iBMDM were washed 3 times with ice cold DPBS, centrifuged at 500 x g, 4°C, for 5 min and then resuspended in ice cold mitochondrial isolation buffer (MIB), composed of 0.25 M sucrose, 20 mM HEPES (pH 7.4), 2 mM EGTA, 10 mM KCl, 1.5 mM MgCl_2_, 0.1% fatty acid-free bovine serum albumin (BSA; A8806, Sigma), and Halt™ Protease inhibitor. Cells were lysed on ice using a sterile 27G syringe and centrifuged at 1100 x g, 4°C, for 3 min to remove unlysed cells and cell debris. Mitochondria were pelleted from the clarified cell lysate by centrifugation at 12,000 x g, 4°C, for 15 min. The supernatant from this step was kept as the cytosolic fraction and the mitochondria-containing pellet (mitochondrial fraction) was resuspended in MIB and centrifuged at 20,000 x g, 4°C, for 10 min. The supernatant was discarded and the mitochondrial fraction was again resuspended in MIB and centrifuged at 20,000 x g, 4°C, for 5 min. The mitochondrial fraction was washed twice with ice cold DPBS at 20,000 x g, 4°C, for 5 min, flash frozen in liquid nitrogen, and stored at −80°C for subsequent analysis.

### Untargeted Lipidomics

WT, CRLS1 KD, and NT-Ctrl macrophages were grown overnight in DMEM supplemented with 10 mM glucose, 1 mM pyruvate, 2 mM glutamine, and 10% FBS. Mitochondria were isolated as described above, flash frozen in liquid nitrogen, and stored at −80°C. Mitochondrial isolates were quality controlled and tested for cytosolic contamination by immunoblot analysis of TOM20 and GAPDH, and total protein stains from SDS-PAGE separated mitochondrial fractions (Revert 700, LI-COR) were used for sample loading normalization for lipidomics data. Untargeted lipidomics analysis was performed after general lipid extraction using a methyl-tert-butyl ether (MTBE)-based liquid-liquid protocol. Samples were thawed at RT and 200 μL of PBS and 500 mL methanol containing 20 mL of an internal standard mixture (custom mixture from Cayman Chemical; see associated documentation) were added to each sample. Samples were vortexed, and 1000 mL methanol and 5 mL MTBE were sequentially added to each sample. After additional vortexing, the mixture was incubated on a tabletop shaker at 500 rpm at RT for 1 hour. Phase separation was induced by the addition of 1.25 mL water. Samples were sonicated for 10 minutes, then centrifuged at 2000 x g for 20 minutes. The upper organic phase of each sample was carefully removed using a Pasteur pipette and transferred into a clean glass tube. The remaining aqueous phase was re-extracted with 2.5 ml of the upper phase of MTBE/methanol/water 10:3:2.5 (v/v/v) solvent mixture, whose composition was similar to the expected composition of the upper phase. After vortexing and centrifugation as above, the organic phase was collected and combined with the initial organic phase. The extracted lipids were dried overnight in a SpeedVac concentrator. The dried lipid extracts were reconstituted in 200 μL n-butanol/methanol 1:1 (v/v) and transferred into autosampler vials for analysis by LC-MS/MS. The parameters of the LC-MS/MS settings can be found in the supplementary data files. Lipostar software (Version 2.0.2b3; Molecular Discovery) was used for feature detection, noise and artifact reduction, alignment, normalization, and lipid identification. Significantly changed lipids between CRLS1 KD and NT-Ctrl macrophages were identified using unpaired T tests and filtering of P < 0.05 and absolute value of log_2_(fold change) > 0.5

### Metabolomics

CRLS1 KD, and NT-Ctrl macrophages were grown overnight in DMEM supplemented with 10 mM glucose, 1 mM pyruvate, 2 mM glutamine, and 10% FBS and then stimulated with or without LPS (200 ng/mL) for 4, 8, or 24h. Stimulation was synchronized such that cells across conditions were in culture for the same amount of time. After stimulation, cells were washed twice with ice cold DPBS, and metabolites were extracted by adding cold 80% methanol, incubating at -80°C for 10 min, followed by centrifugation at 17,000 x g for 10 min at 4°C. The resulting metabolite supernatant was collected. Metabolite extracts were normalized to protein content from paired samples, and the normalized fraction was dried using a SpeedVac at 4°C for 8h. Dried metabolite pellets were resuspended in a 50:50 mixture of MeOH and water. Liquid chromatography-based targeted tandem mass spectrometry (LC-MS/MS)-based metabolomics were performed and the data analyzed as previously described (*46, 47*). In brief, samples were run on an Agilent 1290 Infinity II LC -6470 Triple Quadrupole (QqQ) tandem mass spectrometer system consisting of the 1290 Infinity II LC Flexible Pump (Quaternary Pump), the 1290 Infinity II Multisampler, the 1290 Infinity II Multicolumn Thermostat with 6 port valve and the 6470 triple quad mass spectrometer. Agilent Masshunter Workstation Software LC/MS Data Acquisition for 6400 Series Triple Quadrupole MS with Version B.08.02 was used for compound optimization, calibration, and data acquisition. Significantly changed metabolites between CRLS1 KD and NT-Ctrl macrophages were identified using T tests and filtering of P < 0.05 and absolute value of log_2_(fold change) > 0.5

### Seahorse Extracellular Flux Assay

An Agilent Seahorse XF96 analyzer was used to simultaneously measure the rate of oxygen consumption and extracellular acidification from cultured macrophages. Macrophages were plated in a 96-well Seahorse plate and allowed to adhere overnight in Seahorse XF DMEM supplemented with 10 mM glucose, 1 mM pyruvate, 2 mM glutamine, and 10% FBS. On the following day, media was replaced with or without 200 ng/ml LPS and cells were stimulated for 6h. After 6h, media was exchanged for DMEM supplemented with the same levels of glucose, pyruvate, and glutamine without FBS. Cells were kept in a 37°C incubator without CO_2_ for 30 min prior to analysis. In some experiments, the Mito Stress Test kit from Agilent was used to probe different aspects of mitochondrial function with manufacturer recommended concentrations of respiratory chain inhibitors, 2 μM carbonyl cyanide p-trifluoromethoxyphenylhydrazone (FCCP), 1.5 μM oligomycin, 0.5 μM rotenone, and 0.5 μM antimycin A. Comparable plating between conditions was confirmed by staining plates post analysis with Cell Tracker Red and Hoechst Dye, and analysis using a Biotek Synergy H1 plate reader.

### Protein extraction, SDS-PAGE, and immunoblot (IB) analysis

At experimental endpoints, cells were washed twice with ice-cold DPBS and then lysed in 1% NP40 lysis buffer supplemented with Halt™ Protease and Phosphatase inhibitors for 15 min on ice. A Bio-Rad protein assay was used to normalize sample loading prior to SDS-PAGE. Samples were diluted in Laemmli sample loading buffer, heated for 5 min at 95°C, and then loaded onto precast 4-20% polyacrylamide Tris-Glycine gels (Bio-Rad). After SDS-PAGE, protein was transferred to a 0.45 nitrocellulose membrane by a semi dry transfer system (Cytiva). Membranes were blocked with 5% BSA and 0.1% Tween 20 (IB blocking buffer) for 30 min at RT and then incubated with primary antibody in IB blocking buffer overnight at 4°C. Blots were developed using LI-COR IRdye secondary antibodies and an Odyssey IR Imager. A list of all antibodies used in this study with the associated use and concentration are included in a separate document. Quantification of immunoblots was performed using ImageJ densitometric gel analysis protocol for 1D gels. A list of all antibody sources, concentrations, and applications is available in the associated table.

### Immunoprecipitation (IP) of Complex II

Whole cell extracts were prepared by solubilization in 1% dodecyl maltoside (DDM) supplemented with Halt protease and phosphatase inhibitors on ice for 30 min. The Pierce™ Crosslink immunoprecipitation kit was used to conjugate 5 μg of SDHA antibody (14865-1-AP) to Protein A/G agarose resin. Complex II was immunoprecipitated from whole cell extracts overnight at 4°C with rocking. Complex II was eluted according to the Pierce™ Crosslink immunoprecipitation kit, and the pH was neutralized with pH 9.5 Tris-HCl prior to SDS-PAGE analysis.

### Blue native (BN) PAGE

BN-PAGE analysis of native respiratory complexes was performed as previously described(*48*) using commercially available reagents (Thermo Fisher). In summary, whole cell extracts were prepared by solubilization of 10^6^ cells with 2 mg digitonin and 1X Native PAGE sample buffer (Thermo Fisher, BN2008) on ice for 30 min. Subsequently, insoluble material was removed by 10 min of 17,000g centrifugation at 4°C. The soluble fraction was supplemented with 0.5% Coomassie G-250 immediately before loading samples onto NativePAGE 4 to 16% Bis-Tris mini gels from Invitrogen™. Samples were electrophoresed at 4°C for 30 min in Dark Blue cathode at 150V (Thermo Fisher, BN2007). After this initial electrophoresis, the cathode buffer was switched to the Light Blue cathode buffer. Following electrophoresis, gels were washed with ultrapure water and then soaked with 2x NuPAGE transfer buffer without methanol supplemented with 0.04% SDS for 15 min (Thermo Fisher, NP0006). Protein was transferred to an Immobilon®-FL PVDF membrane at 15V for 15 min with 2x NuPAGE transfer buffer supplemented with 10% methanol using a Bio-Rad Transblot Turbo semi dry transfer system. Immediately following transfer, proteins were fixed in the membrane with 8% acetic acid for 5 min. Samples were washed 3 times with water, 3 times with 100% methanol, and then 3 times with water. Downstream immunoblot analysis was performed normally. Quantification of immunoblots was performed using ImageJ densitometric gel analysis protocol for 1D gels.

### Complex II in gel activity assay

Complex II activity was assessed downstream of native PAGE using an in gel activity assay as previously described(*48*). Briefly, Native PAGE was performed similar to the BN-PAGE protocol above, except that decreased Coomassie G-250 was used to prevent interference with the colorimetric reaction of the activity assay. Following normal BN-PAGE sample loading, samples were electrophoresed at 150V for 30 min at 4°C with the Light Blue Cathode buffer (Thermo Fisher, NP0006). After 30 min, the Light Blue Cathode buffer was replaced with the clear anode buffer and electrophoresis proceeded for an additional 1.5h. Following electrophoresis, the gel was removed and transferred to ice cold ultrapure water. Fresh Complex II activity assay substrate was prepared with the following components: 5mM Tris HCl pH 7.4, 20 mM sodium succinate, 2.5 mg/ml 4-nitroblue tetrazolium, and 2 mM phenazine methosulfate in ultrapure water. 10 mL of the Complex II substrate was incubated on the gel in a 37C incubator for 40 min. The reaction was stopped with 10% acetic acid, washed with ultrapure water, and imaged using a Bio-Rad Gel Doc system.

### Immunofluorescence assay (IFA) and confocal fluorescence microscopic imaging

Macrophages were plated on glass coverslips (No 1.5). Following stimulation, macrophages were fixed at RT with freshly prepared 4% paraformaldehyde in PBS for 15 min. The IFA was always performed on the same day as the experiment. Coverslips were washed three times with PBS + 0.1% Triton-X100 (IFA wash buffer). Coverslips were blocked with a blocking buffer composed of 5% BSA and 10% normal goat serum in wash buffer. A cocktail of primary antibodies was prepared in blocking buffer and incubated on coverslips for 1h at RT. Coverslips were washed three times with IFA wash buffer and incubated with secondary antibodies and counterstains at RT for 30 min. Coverslips were washed and mounted onto microscope slides using Prolong Glass mounting solution. Coverslips were imaged using a Nikon Yokogawa X1-CSU spinning disk confocal microscope equipped with a 100x objective. Operator bias was minimized during image acquisition through selection of fields of view and focal planes based on counterstains which were unrelated to the experimental question. A list of all antibody sources, concentrations, and applications is available in the associated table.

### Confocal microscopic analysis of mitochondrial superoxide (MitoSOX)

Macrophages were plated on glass bottom (No 1.5) Mat-Tek dishes and allowed to adhere overnight. Cells were stimulated +/- LPS for 3 or 6h. Within the last 20 minutes of the experiment, cells were stained with 5 μM MitoSOX dye and 1 μg/ml Hoechst for 20 min at 37°C, protected from light. Cells were washed with PBS and then fixed with 4% PFA at RT for 15 min. Cells were washed with DPBS and then immediately imaged using a Nikon Yokogawa X1-CSU spinning disk confocal microscope. Fields of view were selected based on the Hoechst stain.

### Image analysis and processing

Automated image analysis was performed with the open source software CellProfiler™. All CellProfiler pipelines related to this publication are available upon request. Image analysis was performed on raw images. Automated single cell analysis was achieved by identification of nuclear objects based on global thresholding of a nuclear stain (e.g. Hoechst or DAPI) using the *identify primary objects* module, followed by propagation of the nuclear objects to the cellular periphery based on a whole-cell stain (e.g. Cell Tracker) using the *identify secondary objects* module. Subsequently, a variety of cellular parameters were measured and related to parent cells using the *relate objects* module. **To quantify mitochondrial superoxide**, the intensity of the mitochondrial superoxide dye MitoSOX was measured within cell objects. **To measure SDHB sequestration**, puncta in the SDHB immunostain were identified using the *speckles* enhancement module and Otsu adaptive thresholding within the *identify primary objects* module. Representative images shown in this manuscript were prepared using the ImageJ background subtraction tool with a rolling ball radius of 30 pixels.

### Cytokine analysis

Macrophages were stimulated with or without 200 ng/mL LPS for 6 or 24h. In some experiments, 1 μM or 0.1 μM Atpenin A5 (AA5) or 10 mM, 1 mM, or 0.1 mM of dimethyl malonate (DMM) was added to the culture supernatant for 1h pretreatment. Following stimulation, culture supernatants were collected, and IL-6 and TNF-α levels in the supernatant were measured by ELISA by the University of Michigan Cancer Center Immunology Core.

### Flow cytometric analysis of mitochondrial membrane potential

Macrophages were treated with or without 200 ng/mL LPS for 6h or 20 μM of the protonophore carbonyl cyanide 3-chlorophenylhydrazone (CCCP) for 1h. After stimulation, cells were stained with 150 nM of the mitochondrial membrane potential dye Tetramethylrhodamine methyl ester (TMRM) for 10 min at RT, protected from light. Cells were washed and resuspended in PBS. Cells were immediately analyzed by flow cytometry (Fortessa, BD). Data were processed using FlowJo, and the geometric mean intensity of TMRM from each experiment is reported.

### Real Time quantitative PCR (RT-qPCR)

RNA was extracted from cells using TRIZOL and purified using the Direct-zol RNA MiniPrep Plus kit (Zymo, R2072). The concentration and purity of RNA extracts were determined by Nanodrop. cDNA was synthesized from equivalent amounts of RNA using the iScript cDNA synthesis kit (Bio-Rad, 1708890). RT-qPCR was performed using the SYBR green system and a Bio-Rad CFX96 Real Time system. qPCR primers were designed based on the literature or using NCBI Primer BLAST. A list of all qPCR primers with sources is available in a separate document.

## Acknowledgments

We thank the members of the O’Riordan lab, especially Michael J. McFadden and Zachary M. Powers, for helpful discussions.

## Author contributions

M.B.R., B.H.A., M.X.O. contributed to the conceptualization of the research study. Data curation and formal analysis were completed by M.B.R. and H.S.H. Funding was acquired by M.X.O., C.A.L. and J.S.K. Investigation was completed by M.B.R., H.S.H., B.C.M., L.Z. and A.E.L. Critical resources and supervision were provided by J.S.K., C.A.L. and M.X.O. Writing and visualization of the original manuscript draft were completed by M.B.R, H.S.H. and M.X.O. Review and editing were completed by all authors.

## Data and materials availability

Full lipidomics and metabolomics datasets are available in the supplemental materials of this manuscript. CellProfiler™ image analysis pipelines are available via Github.

## Main Figures

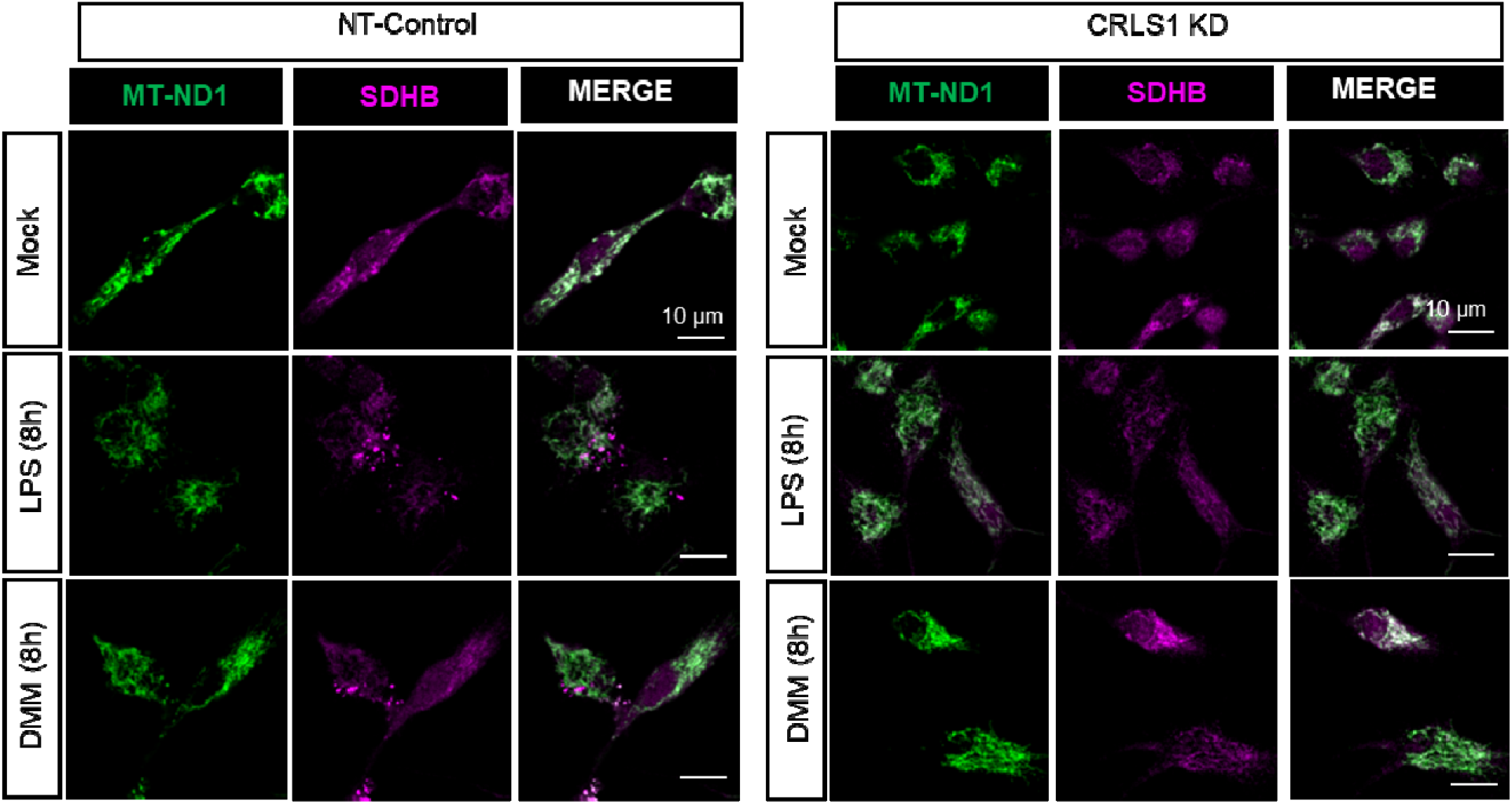

## References

1. L. A. J. O’Neill, R. J. Kishton, J. Rathmell, A guide to immunometabolism for immunologists. Nat. Rev. Immunol. 16, 553–565 (2016).

2. L. Sun (孙李哲), X. Yang (杨晓峰), Z. Yuan (袁祖贻), H. Wang (王虹), Metabolic Reprogramming in Immune Response and Tissue Inflammation. Arterioscler. Thromb. Vasc. Biol. 40, 1990–2001 (2020).

3. A. Viola, F. Munari, R. Sánchez-Rodríguez, T. Scolaro, A. Castegna, The Metabolic Signature of Macrophage Responses. Front. Immunol. 10, 1462 (2019).

4. S. K. Wculek, S. C. Khouili, E. Priego, I. Heras-Murillo, D. Sancho, Metabolic Control of Dendritic Cell Functions: Digesting Information. Frontiers in Immunology. 10 (2019),, doi:10.3389/fimmu.2019.00775.

5. N. M. Chapman, H. Chi, Metabolic adaptation of lymphocytes in immunity and disease. Immunity. 55, 14–30 (2022).

6. B. Kelly, L. A. J. O’Neill, Metabolic reprogramming in macrophages and dendritic cells in innate immunity. Cell Res. 25, 771–784 (2015).

7. Y. Liu, R. Xu, H. Gu, E. Zhang, J. Qu, W. Cao, X. Huang, H. Yan, J. He, Z. Cai, Metabolic reprogramming in macrophage responses. Biomark Res. 9, 1 (2021).

8. A. Chawla, K. D. Nguyen, Y. P. S. Goh, Macrophage-mediated inflammation in metabolic disease. Nat. Rev. Immunol. 11, 738–749 (2011).

9. E. L. Mills, B. Kelly, A. Logan, A. S. H. Costa, M. Varma, C. E. Bryant, P. Tourlomousis, J. H. M. Däbritz, E. Gottlieb, I. Latorre, S. C. Corr, G. McManus, D. Ryan, H. T. Jacobs, M. Szibor, R. J. Xavier, T. Braun, C. Frezza, M. P. Murphy, L. A. O’Neill, Succinate Dehydrogenase Supports Metabolic Repurposing of Mitochondria to Drive Inflammatory Macrophages. Cell. 167, 457-470.e13 (2016).

10. G. M. Tannahill, A. M. Curtis, J. Adamik, E. M. Palsson-McDermott, A. F. McGettrick, G. Goel, C. Frezza, N. J. Bernard, B. Kelly, N. H. Foley, L. Zheng, A. Gardet, Z. Tong, S. S. Jany, S. C. Corr, M. Haneklaus, B. E. Caffrey, K. Pierce, S. Walmsley, F. C. Beasley, E. Cummins, V. Nizet, M. Whyte, C. T. Taylor, H. Lin, S. L. Masters, E. Gottlieb, V. P. Kelly, C. Clish, P. E. Auron, R. J. Xavier, L. A. J. O’Neill, Succinate is an inflammatory signal that induces IL-1β through HIF-1α. Nature. 496, 238–242 (2013).

11. J. K. Dowling, R. Afzal, L. J. Gearing, M. P. Cervantes-Silva, S. Annett, G. M. Davis, C. De Santi, N. Assmann, K. Dettmer, D. J. Gough, G. R. Bantug, F. I. Hamid, F. K. Nally, C. P. Duffy, A. L. Gorman, A. M. Liddicoat, E. C. Lavelle, C. Hess, P. J. Oefner, D. K. Finlay, G. P. Davey, T. Robson, A. M. Curtis, P. J. Hertzog, B. R. G. Williams, C. E. McCoy, Mitochondrial arginase-2 is essential for IL-10 metabolic reprogramming of inflammatory macrophages. Nat. Commun. 12, 1460 (2021).

12. J. Garaude, R. Acín-Pérez, S. Martínez-Cano, M. Enamorado, M. Ugolini, E. Nistal-Villán, S. Hervás-Stubbs, P. Pelegrín, L. E. Sander, J. A. Enríquez, D. Sancho, Mitochondrial respiratory-chain adaptations in macrophages contribute to antibacterial host defense. Nat. Immunol. 17, 1037–1045 (2016).

13. H. Zuo, Y. Wan, Metabolic Reprogramming in Mitochondria of Myeloid Cells. Cells. 9 (2019), doi:10.3390/cells9010005.

14. J. Dudek, Role of Cardiolipin in Mitochondrial Signaling Pathways. Front Cell Dev Biol. 5, 90 (2017).

15. G. Tasseva, H. D. Bai, M. Davidescu, A. Haromy, E. Michelakis, J. E. Vance, Phosphatidylethanolamine deficiency in Mammalian mitochondria impairs oxidative phosphorylation and alters mitochondrial morphology. J. Biol. Chem. 288, 4158–4173 (2013).

16. J. Dudek, M. Hartmann, P. Rehling, The role of mitochondrial cardiolipin in heart function and its implication in cardiac disease. Biochim. Biophys. Acta Mol. Basis Dis. 1865, 810–821 (2019).

17. G. C. Hard, Some biochemical aspects of the immune macrophage. Br. J. Exp. Pathol. 51, 97–105 (1970).

18. P. Newsholme, S. Gordon, E. A. Newsholme, Rates of utilization and fates of glucose, glutamine, pyruvate, fatty acids and ketone bodies by mouse macrophages. Biochem. J. 242, 631–636 (1987).

19. N. V. Dudkina, R. Kouril, K. Peters, H.-P. Braun, E. J. Boekema, Structure and function of mitochondrial supercomplexes. Biochim. Biophys. Acta. 1797, 664–670 (2010).

20. F. Gao, M. B. Reynolds, K. D. Passalacqua, J. Z. Sexton, B. H. Abuaita, M. X. D. O’Riordan, The Mitochondrial Fission Regulator DRP1 Controls Post-Transcriptional Regulation of TNF-α. Front. Cell. Infect. Microbiol. 10, 593805 (2020).

21. V. E. Kagan, J. Jiang, Z. Huang, Y. Y. Tyurina, C. Desbourdes, C. Cottet-Rousselle, H. H. Dar, M. Verma, V. A. Tyurin, A. A. Kapralov, A. Cheikhi, G. Mao, D. Stolz, C. M. St Croix, S. Watkins, Z. Shen, Y. Li, M. L. Greenberg, M. Tokarska-Schlattner, M. Boissan, M.-L. Lacombe, R. M. Epand, C. T. Chu, R. K. Mallampalli, H. Bayır, U. Schlattner, NDPK-D (NM23-H4)-mediated externalization of cardiolipin enables elimination of depolarized mitochondria by mitophagy. Cell Death Differ. 23, 1140–1151 (2016).

22. S. S. Iyer, Q. He, J. R. Janczy, E. I. Elliott, Z. Zhong, A. K. Olivier, J. J. Sadler, V. Knepper-Adrian, R. Han, L. Qiao, S. C. Eisenbarth, W. M. Nauseef, S. L. Cassel, F. S. Sutterwala, Mitochondrial cardiolipin is required for Nlrp3 inflammasome activation. Immunity. 39, 311–323 (2013).

23. Y.-W. Lu, S. M. Claypool, Disorders of phospholipid metabolism: an emerging class of mitochondrial disease due to defects in nuclear genes. Front. Genet. 6, 3 (2015).

24. D. G. Ryan, L. A. J. O’Neill, Krebs Cycle Reborn in Macrophage Immunometabolism. Annu. Rev. Immunol. 38, 289–313 (2020).

25. V. Lampropoulou, A. Sergushichev, M. Bambouskova, S. Nair, E. E. Vincent, E. Loginicheva, L. Cervantes-Barragan, X. Ma, S. C.-C. Huang, T. Griss, C. J. Weinheimer, S. Khader, G. J. Randolph, E. J. Pearce, R. G. Jones, A. Diwan, M. S. Diamond, M. N. Artyomov, Itaconate Links Inhibition of Succinate Dehydrogenase with Macrophage Metabolic Remodeling and Regulation of Inflammation. Cell Metab. 24, 158–166 (2016).

26. W. van der Stel, H. Yang, N. G. Vrijenhoek, J. P. Schimming, G. Callegaro, G. Carta, S. Darici, J. Delp, A. Forsby, A. White, S. le Dévédec, M. Leist, P. Jennings, J. B. Beltman, B. van de Water, E. H. J. Danen, Mapping the cellular response to electron transport chain inhibitors reveals selective signaling networks triggered by mitochondrial perturbation. Arch. Toxicol. 96, 259–285 (2022).

27. M. Yin, L. A. J. O’Neill, The role of the electron transport chain in immunity. FASEB J. 35, e21974 (2021).

28. ezawork-Geleta, H. Wen, L. Dong, B. Yan, J. Vider, S. Boukalova, L. Krobova, K. Vanova, R. Zobalova, M. Sobol, P. Hozak, S. M. Novais, V. Caisova, P. Abaffy, R. Naraine, Y. Pang, T. Zaw, P. Zhang, R. Sindelka, M. Kubista, S. Zuryn, M. P. Molloy, M. V. Berridge, K. Pacak, J. Rohlena, S. Park, J. Neuzil, Alternative assembly of respiratory complex II connects energy stress to metabolic checkpoints. Nat. Commun. 9, 2221 (2018).

29. L. K. Billingham, J. S. Stoolman, K. Vasan, A. E. Rodriguez, T. A. Poor, M. Szibor, H. T. Jacobs, C. R. Reczek, A. Rashidi, P. Zhang, J. Miska, N. S. Chandel, Mitochondrial electron transport chain is necessary for NLRP3 inflammasome activation. Nat. Immunol., 1–13 (2022).

30. S. A. Clayton, K. K. Daley, L. MacDonald, E. Fernandez-Vizarra, G. Bottegoni, J. D. O’Neil, T. Major, D. Griffin, Q. Zhuang, A. B. Adewoye, K. Woolcock, S. W. Jones, C. Goodyear, A. Elmesmari, A. Filer, D. A. Tennant, S. Alivernini, C. D. Buckley, R. D. S. Pitceathly, M. Kurowska-Stolarska, A. R. Clark, Inflammation causes remodeling of mitochondrial cytochrome c oxidase mediated by the bifunctional gene C15orf48. Sci Adv. 7, eabl5182 (2021).

31. B. G. Slane, N. Aykin-Burns, B. J. Smith, A. L. Kalen, P. C. Goswami, F. E. Domann, D. R. Spitz, Mutation of Succinate Dehydrogenase Subunit C Results in Increased O2-, Oxidative Stress, and Genomic Instability. Cancer Res. 66, 7615–7620 (2006).

32. R. D. Guzy, B. Sharma, E. Bell, N. S. Chandel, P. T. Schumacker, Loss of the SdhB, but Not the SdhA, subunit of complex II triggers reactive oxygen species-dependent hypoxia-inducible factor activation and tumorigenesis. Mol. Cell. Biol. 28, 718–731 (2008).

33. B. E. Baysal, R. E. Ferrell, J. E. Willett-Brozick, E. C. Lawrence, D. Myssiorek, A. Bosch, A. van der Mey, P. E. M. Taschner, W. S. Rubinstein, E. N. Myers, C. W. Richard, C. J. Cornelisse, P. Devilee, B. Devlin, Mutations in SDHD, a Mitochondrial Complex II Gene, in Hereditary Paraganglioma. Science. 287, 848–851 (2000).

34. S. Lee, H. Xu, A. Van Vleck, A. M. Mawla, A. M. Li, J. Ye, M. O. Huising, J. P. Annes, β-Cell Succinate Dehydrogenase Deficiency Triggers Metabolic Dysfunction and Insulinopenic Diabetes. Diabetes (2022), doi:10.2337/db21-0834.

35. K. R. Pryde, J. W. Taanman, A. H. Schapira, A LON-ClpP Proteolytic Axis Degrades Complex I to Extinguish ROS Production in Depolarized Mitochondria. Cell Rep. 17, 2522–2531 (2016).

36. I. Martínez-Reyes, N. S. Chandel, Mitochondrial TCA cycle metabolites control physiology and disease. Nat. Commun. 11, 102 (2020).

37. P. Jadiya, D. Tomar, Mitochondrial Protein Quality Control Mechanisms. Genes. 11 (2020), doi:10.3390/genes11050563.

38. R. Kapetanovic, S. F. Afroz, D. Ramnath, G. M. Lawrence, T. Okada, J. E. Curson, J. de Bruin, D. P. Fairlie, K. Schroder, J. C. St John, A. Blumenthal, M. J. Sweet, Lipopolysaccharide promotes Drp1-dependent mitochondrial fission and associated inflammatory responses in macrophages. Immunol. Cell Biol. 98, 528–539 (2020).

39. M. Schlame, Y. Xu, M. Ren, The Basis for Acyl Specificity in the Tafazzin Reaction. J. Biol. Chem. 292, 5499–5506 (2017).

40. O. Ernst, J. Sun, B. Lin, B. Banoth, M. G. Dorrington, J. Liang, B. Schwarz, K. A. Stromberg, S. Katz, S. J. Vayttaden, C. J. Bradfield, N. Slepushkina, C. M. Rice, E. Buehler, J. S. Khillan, D. W. McVicar, C. M. Bosio, C. E. Bryant, F. S. Sutterwala, S. E. Martin, M. Lal-Nag, I. D. C. Fraser, A genome-wide screen uncovers multiple roles for mitochondrial nucleoside diphosphate kinase D in inflammasome activation. Sci. Signal. 14 (2021), doi:10.1126/scisignal.abe0387.

41. S. M. Roberson, W. S. Walker, Immortalization of cloned mouse splenic macrophages with a retrovirus containing the v-raf/mil and v-myc oncogenes. Cell. Immunol. 116, 341–351 (1988).

42. V. Hornung, F. Bauernfeind, A. Halle, E. O. Samstad, H. Kono, K. L. Rock, K. A. Fitzgerald, E. Latz, Silica crystals and aluminum salts activate the NALP3 inflammasome through phagosomal destabilization. Nat. Immunol. 9, 847–856 (2008).

43. D. De Nardo, D. V. Kalvakolanu, E. Latz, Immortalization of Murine Bone Marrow-Derived Macrophages. Methods Mol. Biol. 1784, 35–49 (2018).

44. D. A. Kulpa, N. Del Cid, K. A. Peterson, K. L. Collins, Adaptor protein 1 promotes cross-presentation through the same tyrosine signal in major histocompatibility complex class I as that targeted by HIV-1. J. Virol. 87, 8085–8098 (2013).

45. M. J. Kim, J. W. Hwang, C.-K. Yun, Y. Lee, Y.-S. Choi, Delivery of exogenous mitochondria via centrifugation enhances cellular metabolic function. Sci. Rep. 8, 3330 (2018).

46. C. J. Halbrook, C. Pontious, I. Kovalenko, L. Lapienyte, S. Dreyer, H.-J. Lee, G. Thurston, Y. Zhang, J. Lazarus, P. Sajjakulnukit, H. S. Hong, D. M. Kremer, B. S. Nelson, S. Kemp, L. Zhang, D. Chang, A. Biankin, J. Shi, T. L. Frankel, H. C. Crawford, J. P. Morton, M. Pasca di Magliano, C. A. Lyssiotis, Macrophage-Released Pyrimidines Inhibit Gemcitabine Therapy in Pancreatic Cancer. Cell Metab. 29, 1390-1399.e6 (2019).

47. H.-J. Lee, D. M. Kremer, P. Sajjakulnukit, L. Zhang, C. A. Lyssiotis, A large-scale analysis of targeted metabolomics data from heterogeneous biological samples provides insights into metabolite dynamics. Metabolomics. 15, 103 (2019).

48. P. Jha, X. Wang, J. Auwerx, Analysis of Mitochondrial Respiratory Chain Supercomplexes Using Blue Native Polyacrylamide Gel Electrophoresis (BN-PAGE). Curr. Protoc. Mouse Biol. 6, 1–14 (2016).

