## Supplemental Figures for "Cardiolipin coordinates inflammatory metabolic reprogramming through regulation of Complex II assembly and stability"

**Fig S1 Effect of LPS stimulation on oxygen consumption and extracellular acidification in immortalized bone marrow-derived macrophages (iBMDM).** iBMDM were pretreated with LPS (200 ng/mL) for 6h and then subject to the Agilent Seahorse Extracellular Flux (XF) analysis of the oxygen consumption rate (OCR) and the extracellular acidification rate (ECAR) (**A** and **B**). **C**. Quantification of extracellular lactate as a readout of glycolysis in iBMDM stimulated for 8h with combinations of LPS (200 ng/mL), Complex II inhibitor (Atpenin A5, 1 μM), or vehicle control (0.1% DMSO). Graphs are presented as the mean of n ≥ 3 independent experiments with standard deviation (SD) error bars. P values were calculated using an unpaired T-test or two-way ANOVA with Sidak's post-test for multiple comparisons. *P < 0.05; **P < 0.01; ***P < 0.001; ****P < 0.0001.

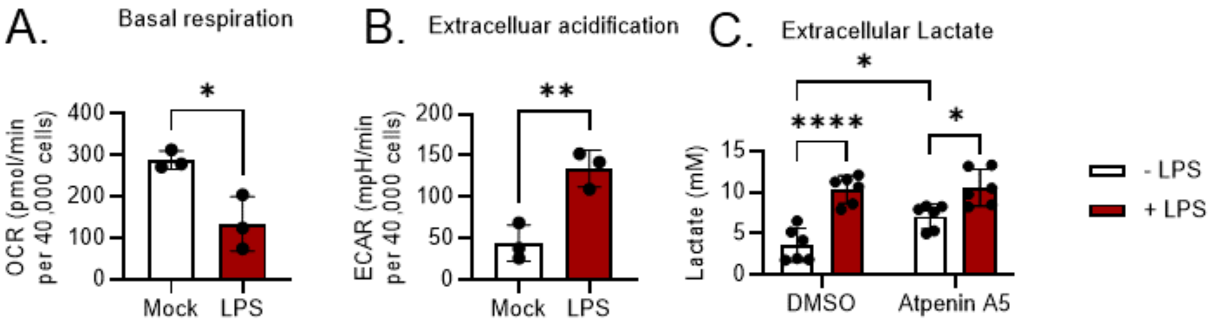

**Fig S2 BN-PAGE analysis of Respiratory Complexes I, II, III, IV, and V during LPS stimulation.** Workflow of parallel native (BN PAGE) respiratory complex immunoblot and Complex II in gel activity assays shown. OXPHOS immunoblot (cocktail of NDUFB8, SDHB, UQCRC2, MTCO1, and ATPA5) and SDHA immunoblot paired to in gel activity assay shown in Fig 1F. As indicated in the analysis workflow, total protein stain (Coomassie) was used as normalization for in gel activity.

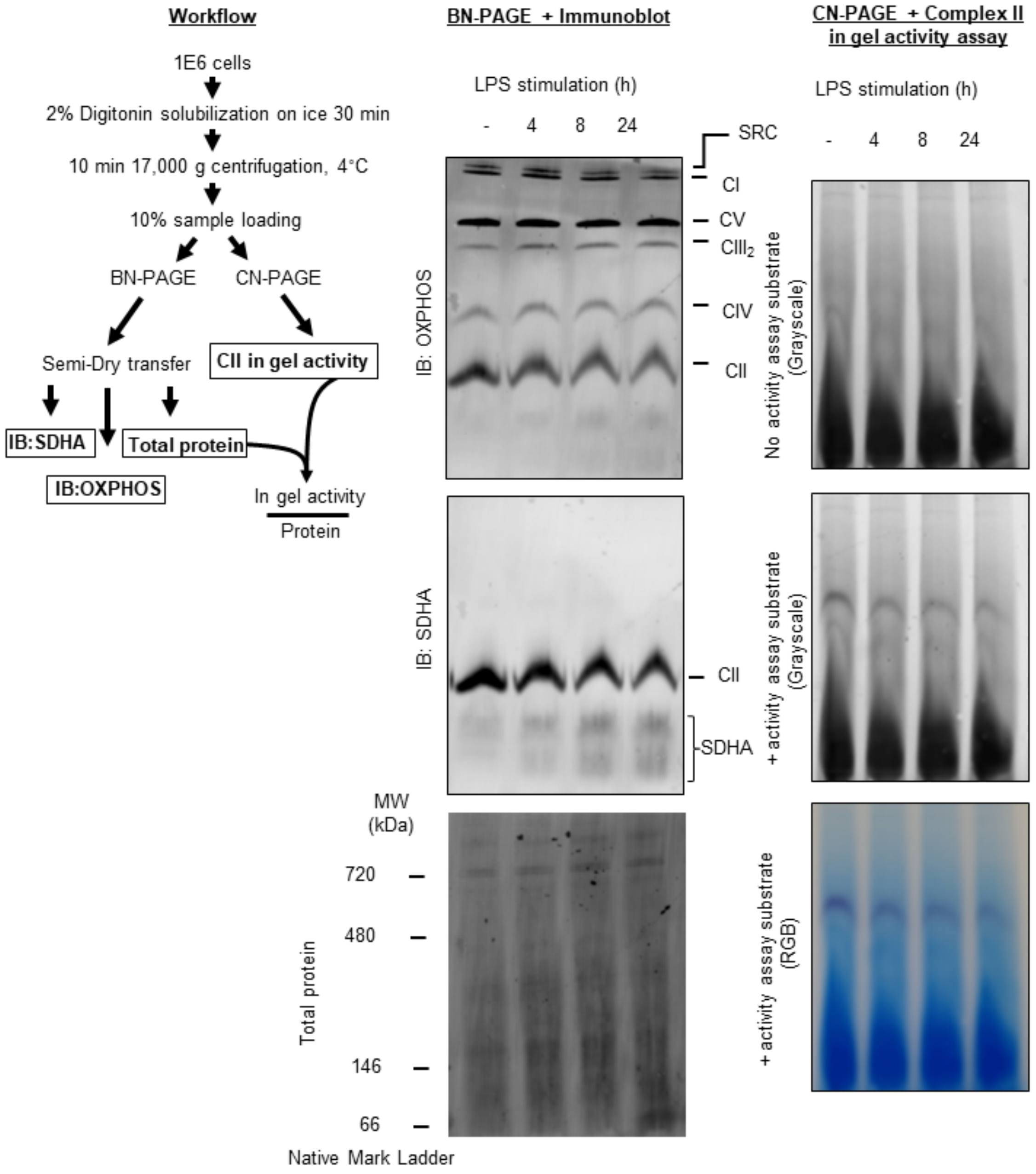

**Fig S3 SDHB and SDHA localization in iBMDM during LPS stimulation.** Representative confocal fluorescence micrographs from iBMDM stimulated with or without 200 ng/mL LPS for 8h and subject to immunofluorescence labeling of Complex II subunit SDHA (**A**), SDHB (**B**), Complex I subunit MT-ND1, and endolysosomal protein LAMP1. Uncropped LAMP1 immunostain from Fig 2B is shown in panel S3B.

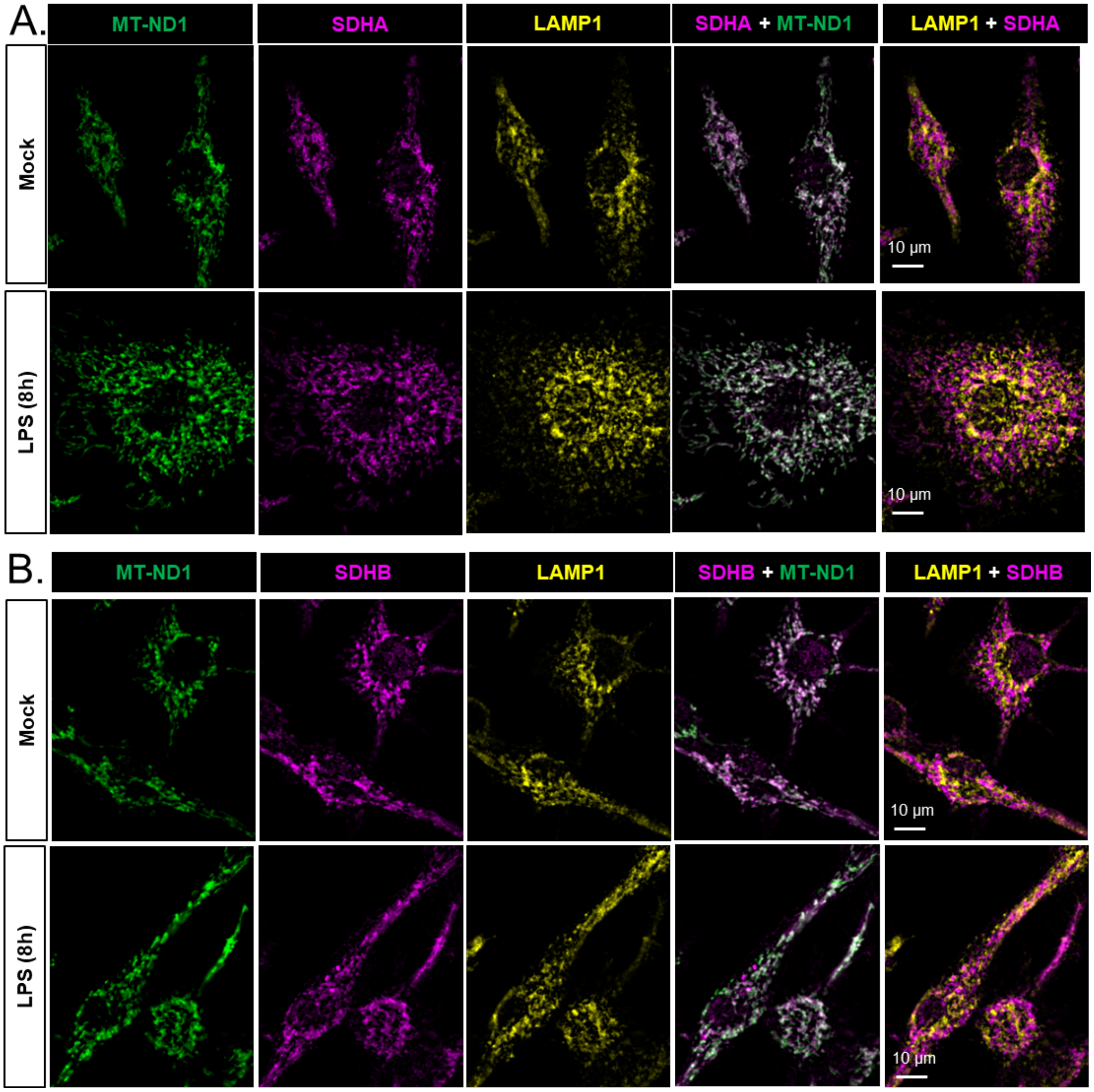

**Fig S4 3D rendering of SDHB puncta and automated image analysis workflow. A.** Representative maximal intensity Z-projections of 10 optical sections, each separated by 0.2 micron, from MT-ND1, LAMP1, and SDHB immunostain in iBMDM stimulated with LPS (200 ng/mL) for 8h. LPS-induced SDHB puncta are emphasized by thresholding the signal of mitochondrial network SDHB (yellow). **B.** 3D render from optical sections described in S4A using Napari and the Ancell Cell and Structure Segmenter plugin. Zoomed regions highlight SDHB puncta localization within the mitochondrial network (MT-ND1 objects) and LAMP1 spherical structures. **C.** Analysis overlay from Cell Profiler of a single optical section of the SDHB immunostain in LPS-treated iBMDM. The white lines indicate the total cell area, while the red lines highlight SDHB puncta identified by the software. **D.** Quantification of the number of SDHB puncta per cell in iBMDM stimulated with or without LPS using the Cell Profiler pipeline described in S4C. Cell-level data pooled from 3 independent experiments and experimental averages are shown. Experimental averages are reported for all of the main figures. Graphs are presented as the mean of $n \geq 3$ independent experiments with standard deviation (SD) error bars. P values were calculated using an unpaired T-test. *$P < 0.05$; ****$P < 0.0001$.

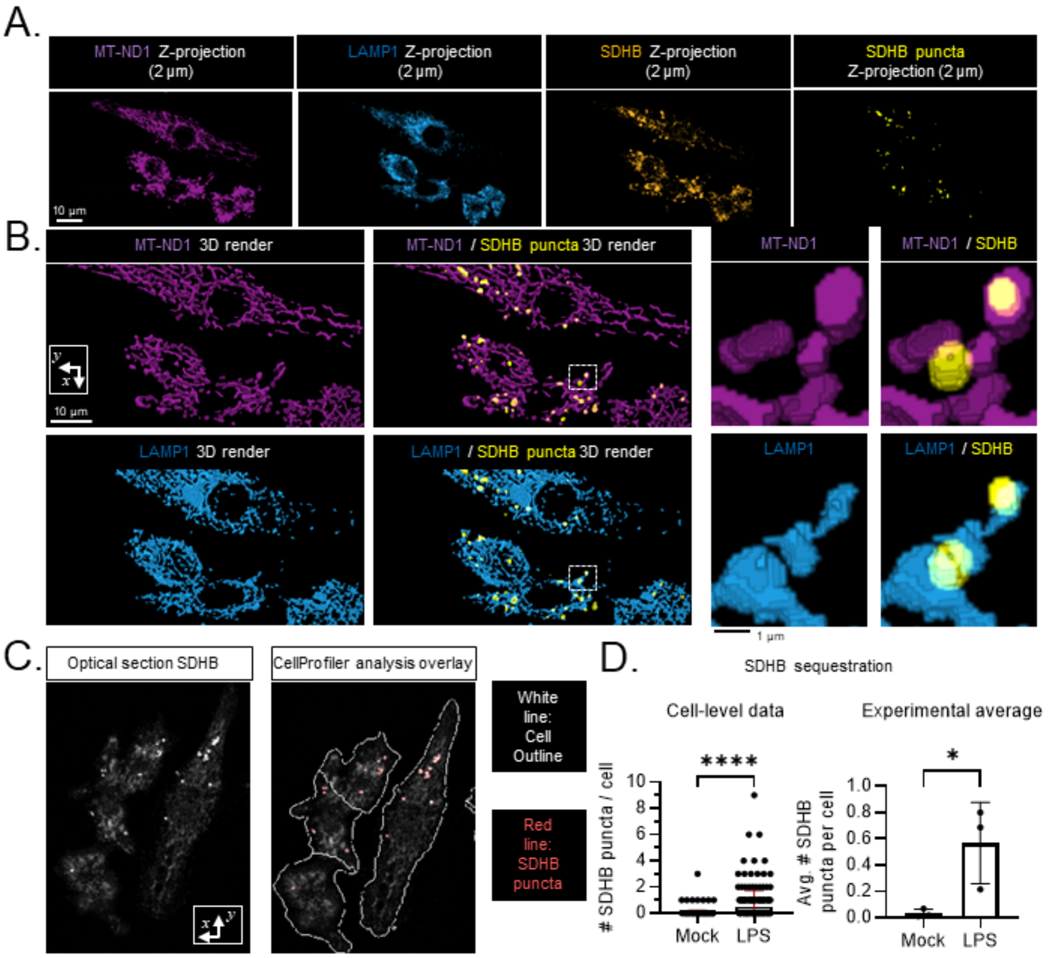
A.
MT-ND1 Z-projection (2 µm)
LAMP1 Z-projection (2 µm)
SDHB Z-projection (2 µm)
SDHB puncta Z-projection (2 µm)
10 µm
B.
MT-ND1 3D render
MT-ND1 / SDHB puncta 3D render
MT-ND1
MT-ND1 / SDHB
10 µm
LAMP1 3D render
LAMP1 / SDHB puncta 3D render
LAMP1
LAMP1 / SDHB
1 µm
C.
Optical section SDHB
CellProfiler analysis overlay
White line: Cell Outline
Red line: SDHB puncta
D.
SDHB sequestration
Cell-level data
Experimental average
### SDHB puncta / cell
Avg. # SDHB puncta per cell
Mock
LPS
****
*

**Fig S5 RT-qPCR analysis of Complex II subunit transcripts *Sdha*, *Sdhb*, *Sdhc*, and *Sdhd*.** The levels of *Sdha*, *Sdhb*, *Sdhc,* and *Sdhd* transcript were measured relative to *Actb* in 6h mock or LPS-treated iBMDM. The fold difference relative to the average mock-treated value for each transcript is reported.

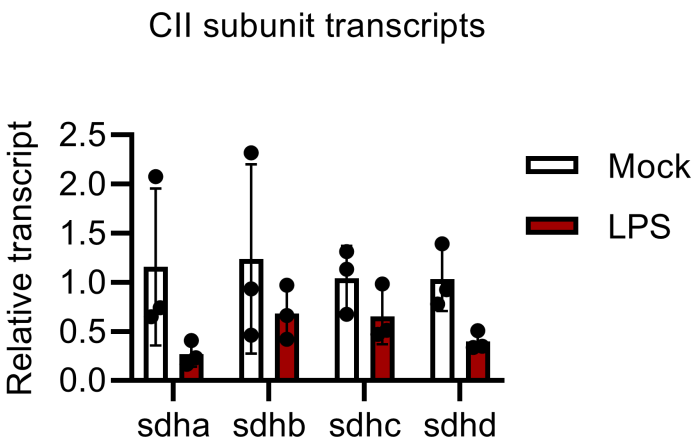

**Fig S6 Untargeted lipidomics analysis of NT-Ctrl and CRLS1 KD iBMDM. A.** Subcellular fractionation of mitochondria and immunoblot analysis of whole cell, cytosolic (GAPDH), and mitochondrial (TOM20) fractions **B.** Significantly changed lipids identified in untargeted lipidomics analysis of mitochondria isolated from CRLS1 KD and NT-Ctrl iBMDM. Peak area is normalized to protein content per sample (Revert700 total protein stain). Lipids levels are reported as $Log_2$(fold change over average level in NT-Ctrl) and sorted by fold change. **C.** All CL species detected in WT, NT-Ctrl, and CRLS1 KD iBMDM are shown. **D.** Table with lipid nomenclature for identified molecular species.

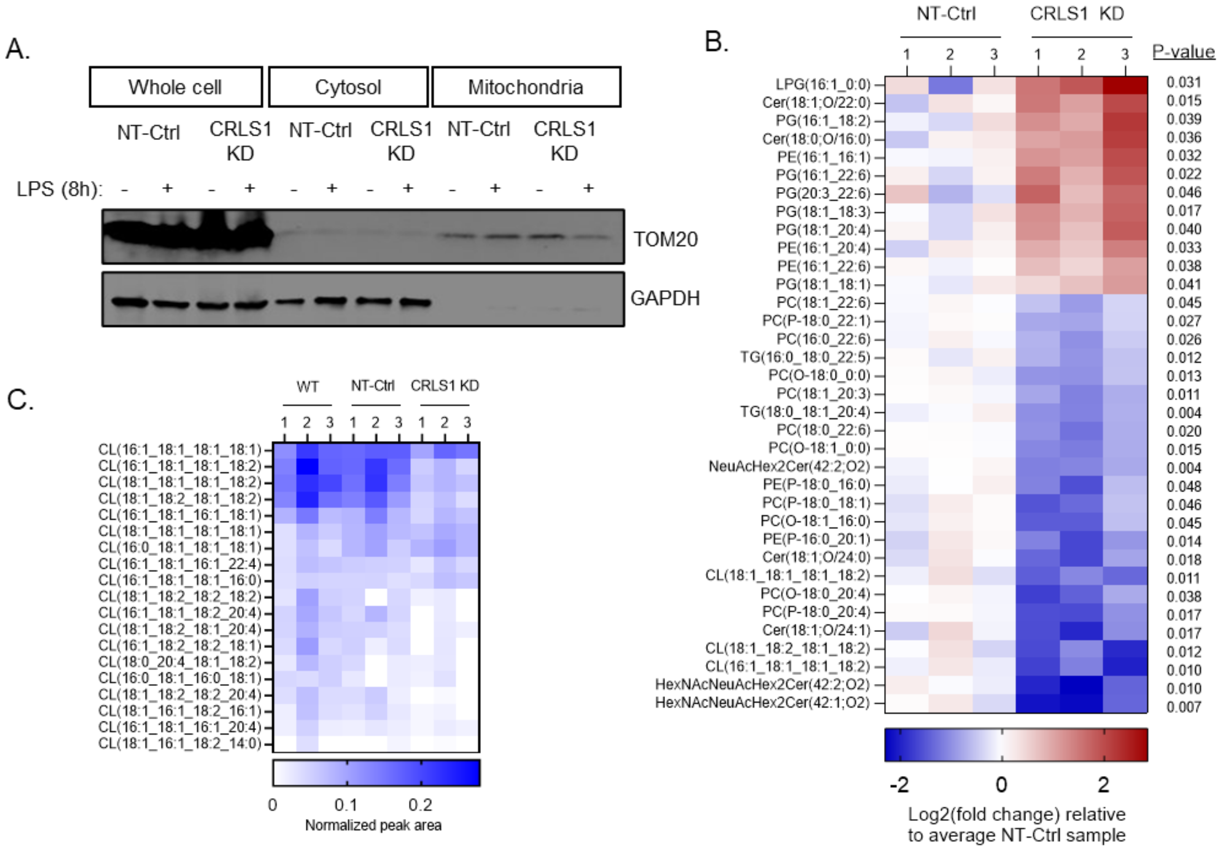


D.

| Molecular Species | | Lipid Class | Lipid Subclass | Category |
|---|---|---|---|---|
| Abbreviated | Common name | | | |
| CL (xx:x_xx:x_xx:x_xx:x) | Cardiolipin | Glycerophosphoglycerophosphoglycerols | Diacylglycerophosphoglycerophosphodiradylglycerols | Glycerophospholipids |
| LPG (xx:x_0:0) | Lysophosphatidylglycerol | Glycerophosphoglycerols | Monoacylglycerophosphoglycerols | Glycerophospholipids |
| PG (xx:x_xx:x) | Phosphatidylglycerol | Glycerophosphoglycerols | Monoacylglycerophosphomonoradylglycerols | Glycerophospholipids |
| PE (xx:x_xx:x) | Phosphatidylethanolamine | Glycerophosphoethanolamines | Diacylglycerophosphoethanolamines | Glycerophospholipids |
| PE (P-xx:x_xx:x) | Phosphatidylethanolamine | Glycerophosphoethanolamines | 1-(1Z-alkenyl),2-acylglycerophosphoethanolamines | Glycerophospholipids |
| PC (xx:x_xx:x) | Phosphatidylcholine | Glycerophosphocholines | Diacylglycerophosphocholines | Glycerophospholipids |
| PC (P-xx:x_xx:x) | Phosphatidylcholine | Glycerophosphocholines | 1-(1Z-alkenyl),2-acylglycerophosphocholines | Glycerophospholipids |
| PC (O-xx:x_xx:x) | Phosphatidylcholine | Glycerophosphocholines | 1-alkyl,2-acylglycerophosphocholines | Glycerophospholipids |
| TG (xx:x_xx:x_xx:x) | Triacylglyceride | Triradylglycerols | Triacylglycerols | Glycerolipids |
| Cer (xx:x;O2/xx:x) | Ceramide | Ceramides | N-acylsphingosines (ceramides) | Sphingolipids |
| NeuAcHex2Cer (xx:x;O2) | Glycosphingolipids | Acidic glycosphingolipids | Gangliosides | Sphingolipids |
| HexNAcNeuAcHex2Cer (xx:x;O2) | Glycosphingolipids | Acidic glycosphingolipids | Gangliosides | Sphingolipids |

**Fig S7 Flow cytometric analysis of TMRM staining in NT-Ctrl and CRLS1 KD iBMDM.** NT-Ctrl and CRLS1 iBMDM were stimulated +/- 200 ng/ml LPS for 6h or 20 μM of the protonophore carbonyl cyanide 3-chlorophenylhydrazone (CCCP) for 1h and then stained with the mitochondrial-membrane potential dye TMRM. 0.1% DMSO was included as a vehicle control for CCCP. The geometric mean intensity of TMRM was measured by flow cytometry per experiment. Representative flow plots are shown. Graphs are presented as the mean of n = 3 independent experiments with standard deviation (SD) error bars. P values were calculated using a two-way ANOVA with Sidak's post-test for multiple comparisons. *P < 0.05 and **P < 0.01.

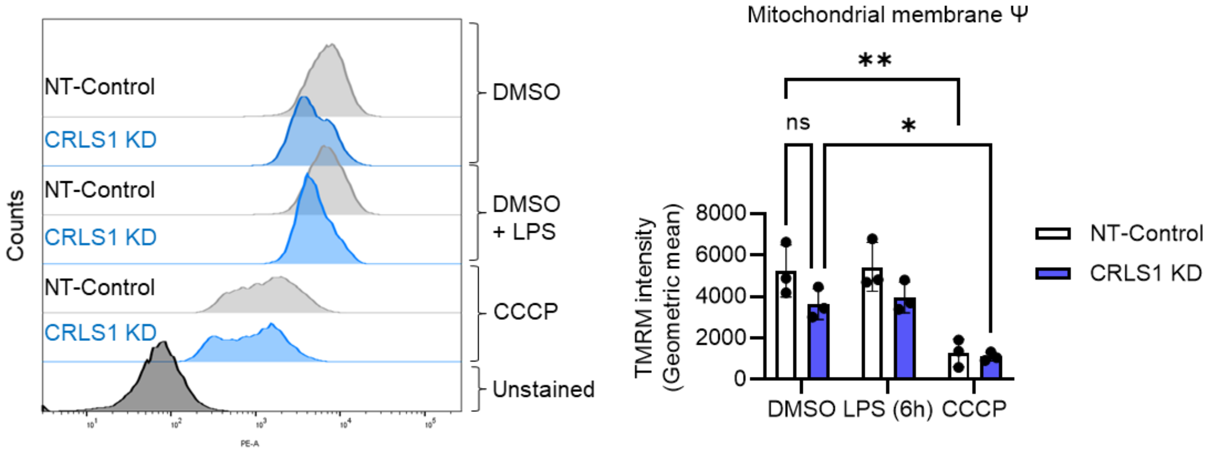

**Fig S8 Targeted metabolomics analysis of NT-Ctrl and CRLS1 KD iBMDM during LPS stimulation.** Significantly changed metabolites identified in targeted metabolomics analysis of whole cell extracts from CRLS1 KD and NT-Ctrl iBMDM stimulated +/- 200 ng/ml LPS for 4, 8, or 24h. Metabolite levels are reported as $Log_2$(fold change over average level in unstimulated NT-Ctrl). Metabolites are sorted by P-value following T test comparison of 24h LPS-treated NT-Ctrl and CRLS1 KD iBMDM. Identified TCA-related metabolites are bolded and notated with an asterisk.

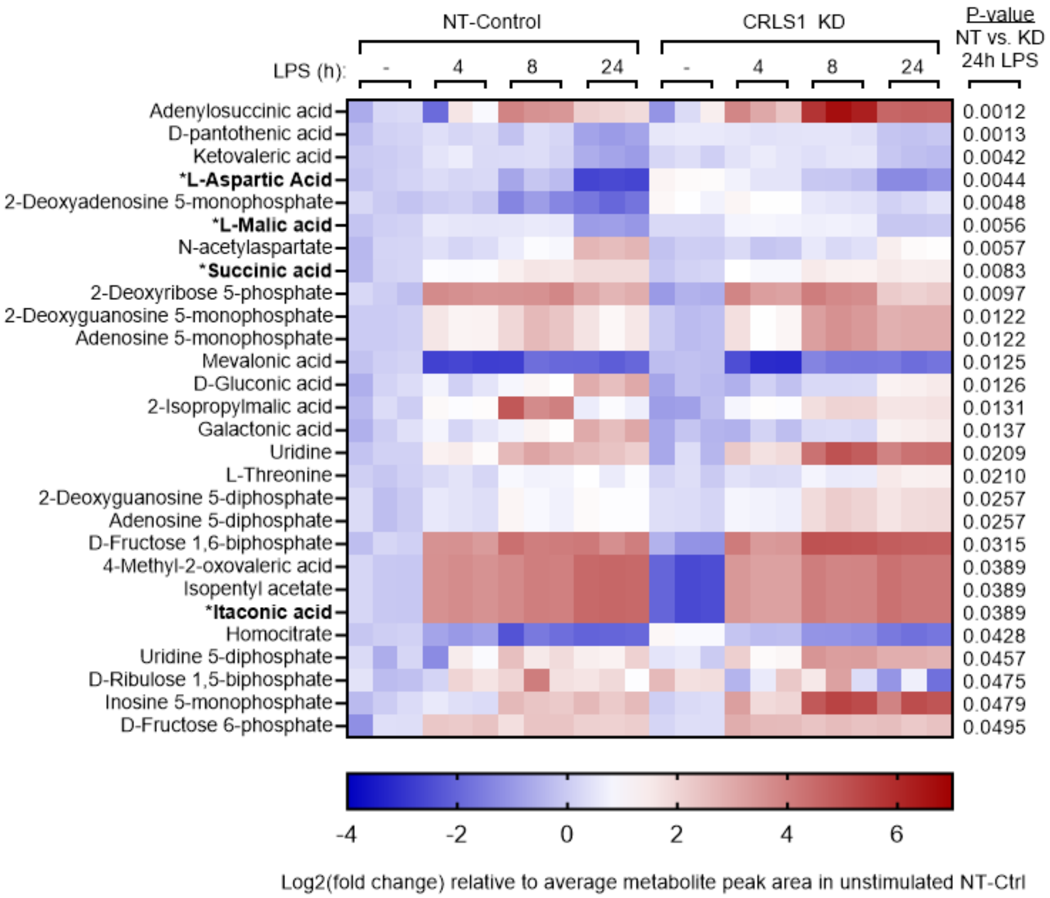

**Fig S9 Proinflammatory cytokine production in LPS-stimulated NT-Ctrl and CRLS1 KD RAW264.7 cells.** RAW264.7 cells were transduced with a lentiviral vector containing an shRNA targeted to *Crls1* or an non-target control (NT-Ctrl) sequence. **A.** *Crls1* levels in CRLS1 KD and NT-Ctrl RAW264.7 cells. **B.** Transcript levels of *Il6 and Tnf* in CRLS1 KD and NT-Ctrl RAW264.7 cells stimulated +/- 200 ng/ml LPS for 4h. **C.** ELISA analysis of secreted IL-6 and TNF-α in CRLS1 KD and NT-Ctrl RAW264.7 cell stimulated +/- 200 ng/ml LPS for 24h. Graphs are presented as the mean of n = 3 independent experiments with standard deviation (SD) error bars. P values were calculated using a two-way ANOVA with Sidak's post-test for multiple comparisons. *P < 0.05; **P < 0.01; ***P < 0.001; and ****P < 0.0001.

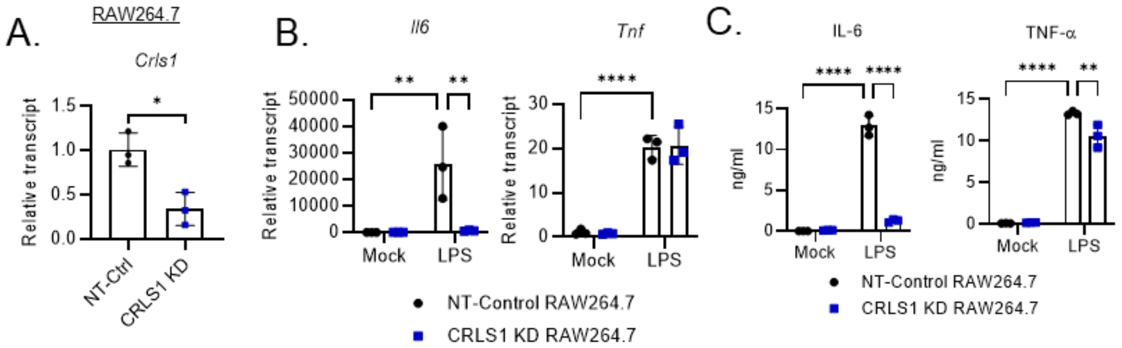

**Fig S10 IL-6 secretion in *Salmonella enterica* Typhimurium-infected NT-Ctrl and CRLS1 KD iBMDM.** ELISA analysis of secreted IL-6 in CRLS1 KD and NT-Ctrl iBMDM infected with MOI 20 of *Salmonella enterica* serovar Typhimurium (STM) (**SL1344**) for 24h. Graphs are presented as the mean of n = 3 independent experiments with standard deviation (SD) error bars. P values were calculated using a two-way ANOVA with Sidak's post-test for multiple comparisons. **P < 0.01.

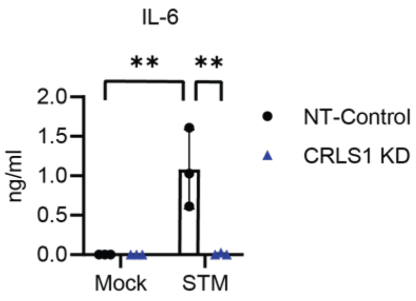

**Fig S11 IL-6 and TNF-α secretion CRLS1 KD with Atpenin A5 treatment.** ELISA analysis of IL-6 and TNF-α secreted by NT-Ctrl and CRLS1 KD iBMDM pretreated for 1h with or without 0.1 or 1 uM Atpenin A5 (AA5) then challenged with 200 ng/mL LPS for 6h. 0.1% DMSO was included as a vehicle control for AA5. Graphs are presented as the mean of n ≥ 3 independent experiments with standard deviation (SD) error bars. P values were calculated using a two-way ANOVA with Sidak's post-test for multiple comparisons. *P < 0.05; **P < 0.01; ***P < 0.001; and ****P < 0.0001.

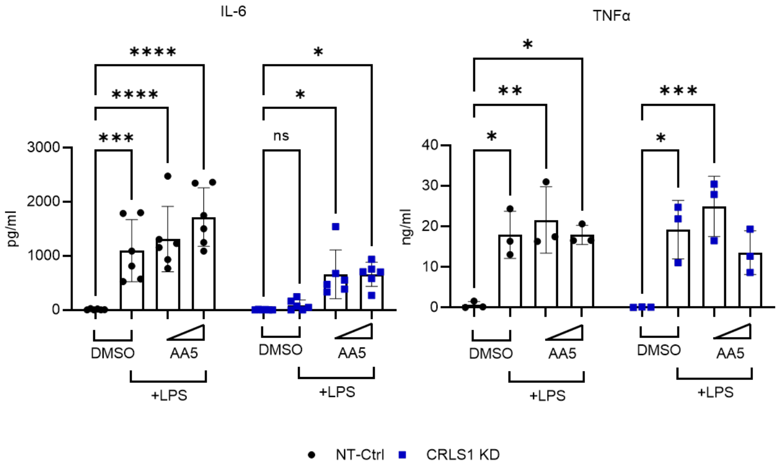

**Fig S12 Exogenous delivery of diethyl succinate (DES) does not restore CRLS1-dependent IL-6 production.** CRLS1 KD and NT-Ctrl iBMDM were pre-treated with 5 mM diethyl succinate (DES) for 2h and then challenged with 200 ng/mL LPS for 6 h. Supernatants were collected and secreted IL-6 was analyzed by ELISA. Graphs are presented as the mean of n = 3 independent experiments with standard deviation (SD) error bars. P values were calculated using a two-way ANOVA with Sidak's post-test for multiple comparisons. *P < 0.05 and **P < 0.01

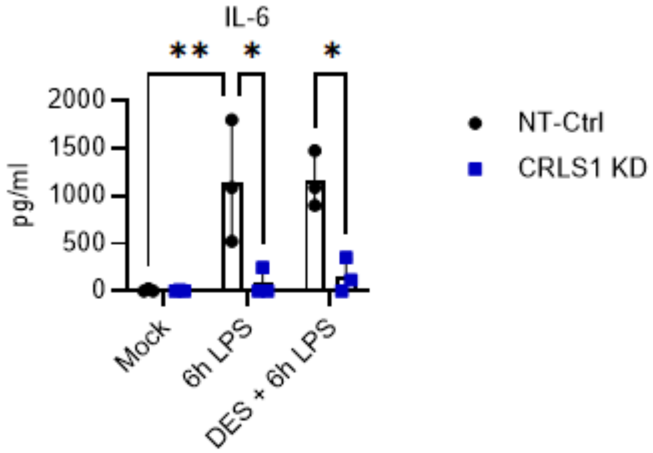

**Fig S13 SDHB sequestration during dimethyl malonate (DMM) treatment.** Representative confocal fluorescence micrographs from NT-Ctrl and CRLS1 KD iBMDM stimulated with or without 10 mM DMM or 200 ng/ml LPS for 8h and subject to immunofluorescence labeling of SDHB and MT-ND1.

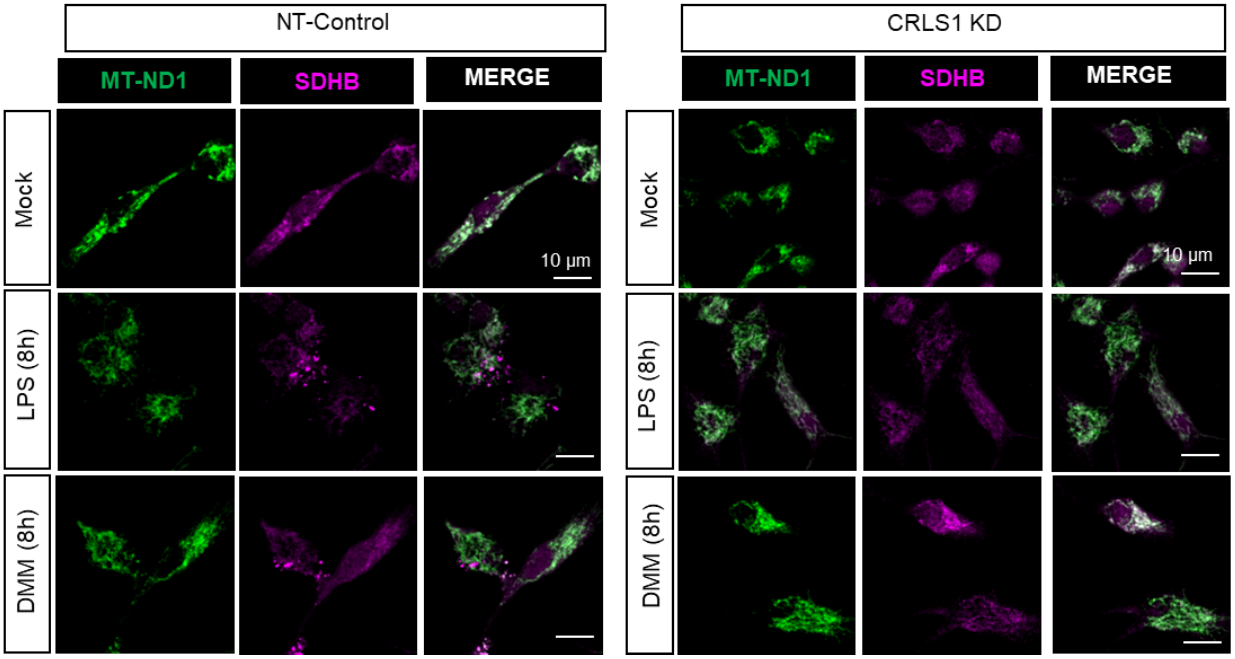
